# Lesion-specific suppression of YAP/TAZ by biomimetic nanodrug ameliorates atherosclerosis development

**DOI:** 10.1101/2023.04.24.537992

**Authors:** Hui-Chun Huang, Ting-Yun Wang, Joshua Rousseau, Michelle Mungaray, Chamonix Michaud, Christopher Plaisier, Zhen Bouman Chen, Kuei-Chun Wang

## Abstract

Atherosclerosis, characterized by the buildup of lipid-rich plaque on the vessel wall, is the primary cause of myocardial infarction and ischemic stroke. Recent studies have demonstrated that dysregulation of yes-associated protein 1 (YAP) and transcriptional coactivator with PDZ-binding domain (TAZ) contributes to plaque development, making YAP/TAZ potential therapeutic targets. However, systemic modulation of YAP/TAZ expression or activities risks serious off-target effects, limiting clinical applicability. To address the challenge, this study develops monocyte membrane-coated nanoparticles (MoNP) as a drug delivery vehicle targeting activated endothelium lining the plaque surface and utilizes MoNP to deliver verteporfin (VP), a potent YAP/TAZ inhibitor, for lesion-specific treatment of atherosclerosis. The results reveal that MoNP significantly enhance payload delivery to inflamed endothelial cells (EC) while avoiding phagocytic cells, and preferentially accumulate in atherosclerotic regions. MoNP-mediated delivery of VP substantially reduces YAP/TAZ expression, suppressing inflammatory gene expression and macrophage infiltration in cultured EC and mouse arteries exposed to atherogenic stimuli. Importantly, this lesion-targeted VP nanodrug effectively decreases plaque development in mice without causing noticeable histopathological changes in major organs. Collectively, these findings demonstrate a plaque-targeted and pathway-specific biomimetic nanodrug, potentially leading to safer and more effective treatments for atherosclerosis.

## 1. Introduction

Maladaptive inflammation, a key feature in all stages of atherosclerosis, exacerbates plaque development and progression, prompting scientists to explore the potential of anti-inflammatory therapies for atherosclerosis [1,2]. Recent clinical trials have demonstrated the feasibility of anti-inflammatory agents in reducing secondary cardiovascular events, positioning them as potential future therapeutic options for atherosclerosis, alongside statin treatments [3,4]. Despite the success, the impact of systemic cytokine neutralization on plaque characteristics, as well as issues associated with impaired host defense and infections, remain to be addressed [5,6]. In comparison to systemic treatments, local delivery of therapeutics that directly target specific inflammatory pathways in diseased blood vessels would be a more preferable approach to treat plaque progression, potentially improving effectiveness and safety. However, the complex nature of plaque makes the development of a lesion-targeted, pathway-specific strategy a highly challenging goal.

Emerging evidence, including ours, revealed that many of the pathological events in atherosclerosis are mediated by dysregulation of yes-associated protein 1 (YAP) and transcriptional coactivator with PDZ-binding domain (TAZ) [7–9]. YAP/TAZ are primary effectors of the Hippo pathway, and upon activation, YAP/TAZ translocate into the nucleus and interact with transcriptional enhanced associate domain (TEAD) proteins to promote the expression of specific genes, including connective tissue growth factor (CTGF) and those involved in inflammation [10,11]. In response to atheroprone stimuli, such as hemodynamic disturbances and proinflammatory cytokines, the activities of YAP/TAZ in vascular endothelial cells (EC) are greatly increased, promoting macrophage infiltration into the arterial wall and plaque development [7,9,12]. More importantly, we and others have demonstrated that inhibiting the expression of YAP and/or TAZ reduced arterial inflammation and plaque size in hyperlipidemic mice, suggesting YAP/TAZ are promising therapeutic targets for atherosclerosis [7,8].

Verteporfin (VP) is an FDA-approved, light-activated agent for age-related macular degeneration [13]. In addition to its function as a photosensitizer, VP has been shown to reduce YAP/TAZ expression and block the interactions between YAP and TEAD proteins in the absence of photoactivation, making VP a promising inhibitor of YAP/TAZ functions [14,15]. Given its potency in YAP/TAZ inhibition, VP has been repositioned as a light-independent agent to treat YAP/TAZ-induced pathologies, such as cancers, in numerous preclinical models and a clinical trial [16,17]. Despite its potential, our knowledge of VP treatment in atherosclerosis and other inflammatory diseases involving YAP/TAZ dysregulation remains very limited. In this study, we sought to investigate whether VP treatment can mitigate atherosclerosis through a light-independent reaction, specifically by suppressing YAP/TAZ activities in inflamed endothelium. However, systemic administration of VP may cause YAP/TAZ inhibition beyond the atherosclerotic vasculature, raising concerns about indirect and undesired side effects. Thus, a lesion-targeted VP nanodrug that enables local inhibition of YAP/TAZ in diseased arteries, minimizing unintended delivery to healthy vessels, would be highly desirable.

Cell membrane-coated nanoparticles (NP), leveraging unique biomimetic features from the source cells to enhance the biocompatibility, stealth ability, and functionality, present an appealing drug delivery platform and have demonstrated promising results in various disease models [18,19]. For instance, NP coated with red blood cell membrane have demonstrated the capability of immune evasion, extending their circulation time compared to uncoated NP [20]. Similarly, cancer cell membrane-cloaked NP have been engineered to attain homotypic targeting to tumor tissues [21]. In the context of cardiovascular disease, NP harnessing the membrane of platelets or immune cells improved drug localization to injured vessels, plaques, and ischemic hearts, respectively, thereby enhancing therapeutic outcomes [22–24]. Building on these successes, this present study aimed to develop a biomimetic drug delivery platform for active targeting of intimal endothelium lining atherosclerotic arteries. Considering that the influx of monocytes (Mo)/macrophages to the arterial wall is a critical feature in all stages of atherosclerosis, we explored the potential of using plasma membrane from primary Mo to functionalize NP. The rationale is that Mo membrane-coated NP (MoNP), with the inherent capabilities of classical Mo, could evade clearance by the mononuclear phagocyte system (MPS) and adhere to inflamed endothelium on the plaque surface. By employing MoNP as a targeted delivery vehicle, we tested the hypothesis that selective delivery of VP to atherosclerotic sites could be achieved, further allowing us to examine the therapeutic efficacy of this localized anti-YAP/TAZ therapy in atherosclerosis.

In this study, we demonstrated that Mo membrane cloaking strongly increased the uptake of NP by inflamed EC, but not phagocytic cells, and promoted their accumulation in inflamed vasculature rather than healthy vessels, confirming the enhanced ability of MoNP for selective targeting and immune evasion. By utilizing MoNP to deliver VP, we observed significant reductions in arterial YAP/TAZ expression, EC inflammation, macrophage infiltration, and plaque formation using *in vitro* and *in vivo* models without causing significant organ toxicity in mice. Collectively, our findings support that MoNP-VP treatment holds great potential as a safe and effective strategy for treating atherosclerosis.

## 2. Results

### 2.1. Formulation and characterization of MoNP

To achieve active targeting to inflamed endothelium and plaques of the vasculature, we first prepared the biomimetic MoNP using the plasma membrane of mouse Mo and poly(lactic-co-glycolic acid) (PLGA) NP (Figure 1A). Briefly, NP were prepared using an oil-in-water-emulsion method, after which they were cloaked with the membrane vesicles from mouse classical Mo (Ly6c^+^/CD11b^+^/CCR2^+^/CD45R^-^) isolated from the suspension of bone marrow cells (BMC) differentiated in the medium containing M-CSF (Figure 1B and S1). The resulting MoNP were then subjected to physicochemical characterization using dynamic light scattering (DLS), transmission electron microscopy (TEM), SDS-PAGE, and Western blot analyses. Our DLS data showed that MoNP were highly uniform in size, with an average hydrodynamic size of 284.74 nm, while Mo vesicles and uncoated NP are 430.26 nm and 216.36 nm, respectively (Figure 1C). MoNP had a mean surface charge of -17.45 mV, which was much closer to that of Mo vesicles than NP (i.e., -11.88 mV for Mo vesicles and -31.26 mV for NP), indicating successful membrane cloaking (Figure 1D). The TEM analysis confirmed that Mo membrane was fused onto NP to form a core-shell nanostructure with a final diameter slightly over 200 nm (Figure 1E). Furthermore, our SDS-PAGE and Western blot results demonstrated that CD11b, Na^+^/K^+^ ATPase, and TLR4, but not GAPDH, were decorated on the surface of MoNP, verifying that Mo membrane proteins, rather than their intracellular components, were present on MoNP (Figure 1F and 1G).

**Figure 1.**
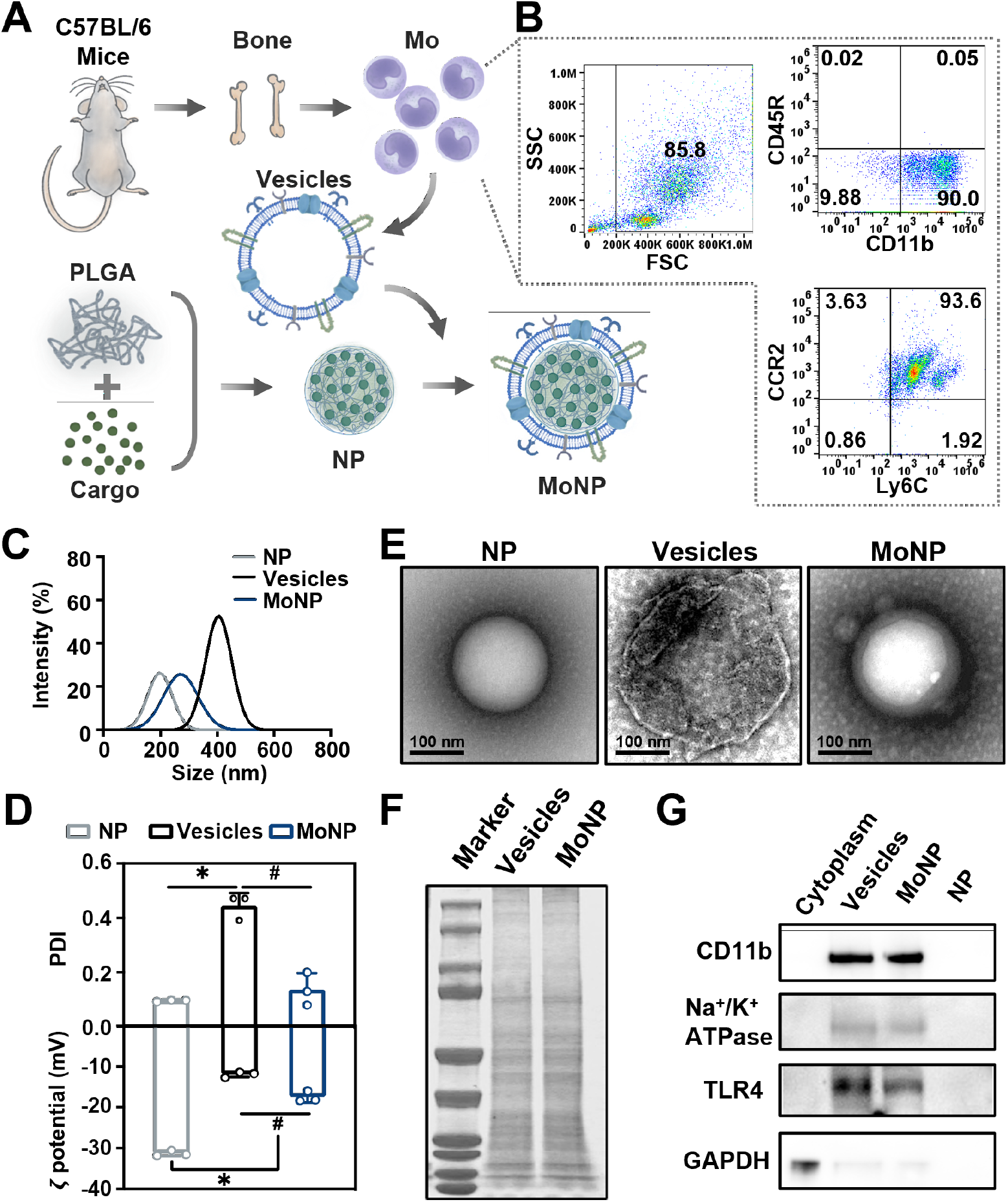
Formulation and characterization of MoNP. (A) A schematic illustrating the preparation of MoNP loaded with a therapeutic payload. (B) Flow cytometric analysis of Mo isolated from differentiated mouse BMC. (C) and (D) DLS results of MoNP, NP, and Mo vesicles; (C) hydrodynamic size, (D) polydispersity index (PDI), and surface charge. (E) Representative TEM images of MoNP, NP, and Mo vesicles. Scale bar = 100 nm. (F) SDS-PAGE analysis of membrane protein profiles of MoNP and Mo vesicles. (G) Western blot analysis of the membrane proteins of MoNP and Mo vesicles. Graphic data in (D): n = 3, *p < 0.05 vs. NP and ^#^p < 0.05 vs. Mo vesicles.

We next investigated the stability of MoNP by storing aliquots at different temperatures (25°C, 4°C, or -20°C) for 4 days. Our findings showed that storing MoNP at -20°C significantly reduced particle aggregation and helped maintain monodispersity even after the freezing-thawing process (Figure S2). To assess the drug loading capacity and release profile, we formulated MoNP with the fluorescent dye DiD as a payload. Approximately 61% of the input DiD could be encapsulated into MoNP, attaining a loading efficiency (LE) of 3.3%, and the final hydrodynamic size remained stable at approximately 290 nm (Figure S3). Additionally, when incubated with a saline solution containing 10% serum, approximately 90% of the payload could be released from NP within 24 hours (Figure S4).

We next assessed the biocompatibility of the MoNP by evaluating their *in vitro* and *in vivo* toxicity. First, we incubated EC with MoNP and monitored their growth, and we observed that the presence of MoNP did not affect cell proliferation (Figure S5A). We also incubated mouse whole blood with MoNP and found that MoNP did not cause apparent hemolysis (Figure S5B). Moreover, we intravenously administered MoNP to mice and evaluated organ toxicity via histological examination. MoNP did not induce any noticeable histopathological changes in major organs, including the heart, liver, kidney, lung, and spleen (Figure S5C). These results indicate that MoNP is a highly biocompatible vehicle for drug delivery, with no significant toxic effects observed in cellular or animal models.

### 2.2. MoNP facilitates selective uptake by inflamed EC rather than phagocytes

To investigate the impact of MoNP on cellular uptake, we incubated EC and Mo, respectively, with MoNP-DiD or uncoated NP-DiD, followed by analyses of the intracellular DiD signal using fluorescence microscopy and flow cytometry. MoNP-DiD treatment, compared to uncoated NP-DiD, significantly increased the number of DiD-positive EC, especially in those pre-treated with proinflammatory cytokine TNFα. Notably, pretreatment of an antibody against VCAM1 abolished the increased uptake of MoNP-DiD in inflamed EC (Figure 2A and S6A). We also tested the uptake of MoNP-DiD by EC exposed to different magnitudes of shear stress (SS) [7]. Using a parallel plate flow system, we observed more MoNP accumulated in EC under low SS, which is commonly observed in atheroprone regions and found to promote EC inflammation, than those under high SS (Figure 2B and S6B). In contrast to the findings in EC, mouse phagocytes, including Mo and macrophages, internalized more uncoated particles than those enclosed in Mo membrane (Figure 2C, 2D, S6C, and S6D). Additionally, we incubated mouse whole blood with MoNP-DiD or NP-DiD and found a stronger DiD signal in the plasma of the MoNP-DiD group than that of NP-DiD (Figure S6E). These results suggest that Mo membrane cloaking enhances the interaction between MoNP and inflamed endothelium in a VCAM1-dependent manner and reduces uptake by immune phagocytes. To investigate the intracellular fate of MoNP, we incubated EC with MoNP-DiD or NP-DiD and then stained lysosomes using LysoTracker Green. Compared to NP, MoNP showed a lower colocalization with lysosomes than the NP group, indicating that Mo membrane coating could facilitate the escape of resulting particles from lysosomes after internalization and potentially extend their therapeutic effects (Figure 2E and 2F).

**Figure 2.**
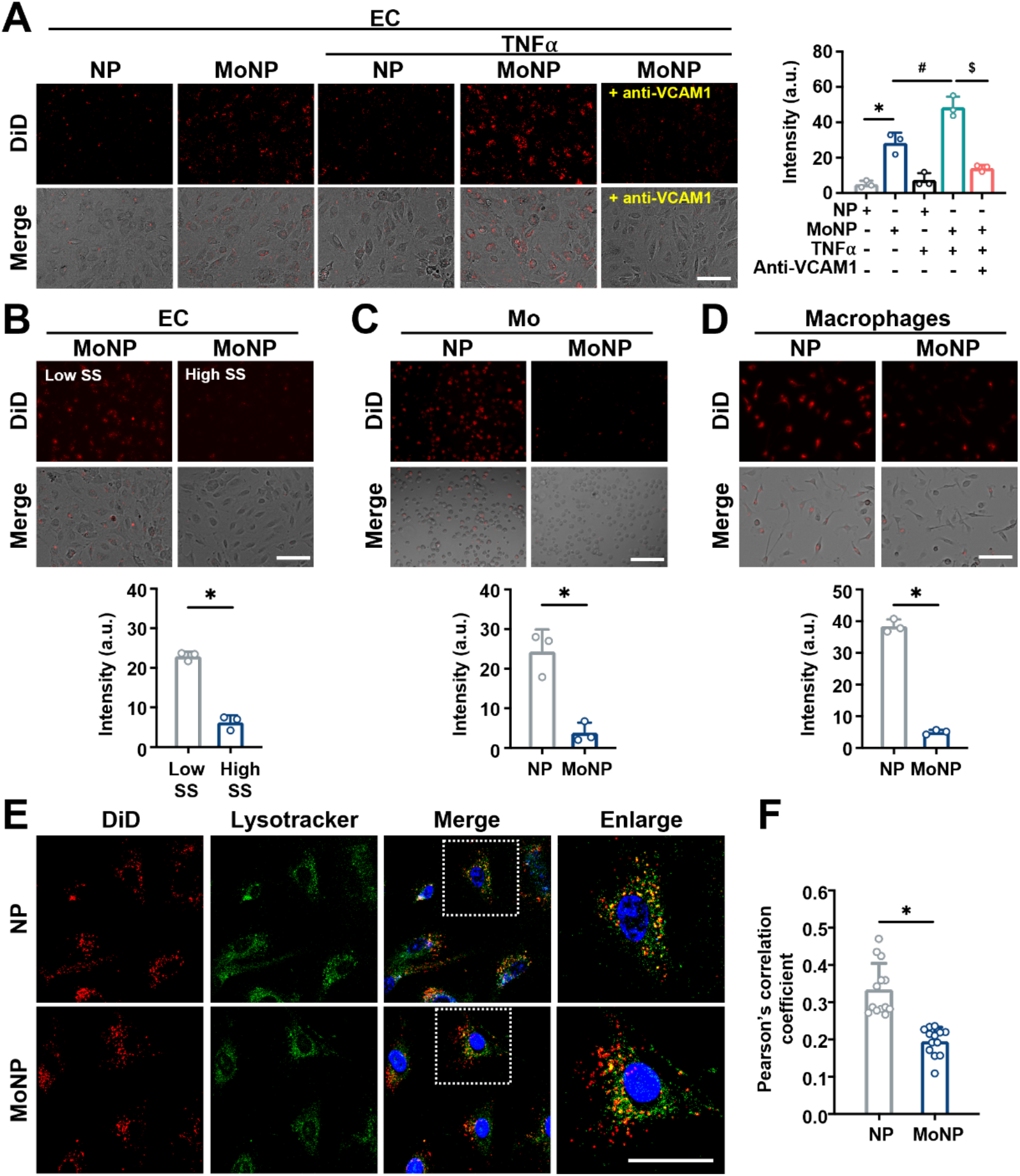
MoNP enhanced endothelial uptake and lysosomal escape. (A-C) Representative fluorescent images showing (A) the cellular uptake of MoNP-DiD or NP-DiD by EC, TNFα-pretreated EC, TNFα-/anti-VCAM1-pretreated EC, (B) the cellular uptake of MoNP-DiD by EC under low or high SS, and (C-D) the cellular uptake of MoNP-DiD or NP-DiD by (C) Mo and (D) macrophages. Scale bar = 100 μm. The intracellular DiD signal was quantified. For A, C, and D, n = 3; *p < 0.05 vs. NP-DiD, ^#^p < 0.05 vs. MoNP-DiD, ^$^p < 0.05 vs. MoNP-DiD/TNFα-stimulated EC. For B, n = 3; *p < 0.05 vs. low SS. (E) Representative confocal images of EC incubated with MoNP-DiD or NP-DiD (red), followed by the staining of lysotracker (green) and nuclei (blue). Scale bar = 25 μm. (F) Correlation analysis of MoNP-DiD or NP-DiD with lysosomes. Thirteen cells were randomly selected from fluorescent images acquired in three biological repeats. *p < 0.05 vs. NP-DiD.

### 2.3. MoNP preferentially accumulates in the intima of atherosclerotic arteries

To determine whether MoNP could achieve active targeting of inflamed EC lining atherosclerotic vasculature *in vivo*, we performed the partial ligation (PL) procedure in apolipoprotein E-deficient (ApoE^-/-^) mice. PL creates flow disturbance in the left carotid artery (LCA) and is a well-established approach to induce endothelial activation/dysfunction and accelerate the development of carotid atherosclerosis [25]. Seven days after PL, we intravenously administered MoNP-DiD or NP-DiD to the mice. Four hours post-injection, we harvested arterial tissues from descending aorta to carotid bifurcations and major organs to determine the biodistribution of administered particles (Figure 3A). Fluorescent imaging of whole-mount arterial tissues revealed a strong DiD signal in the partially ligated LCA, followed by aortic arches, in mice receiving MoNP-DiD but not in those receiving NP-DiD. Additionally, only background signal was detected in intact regions of the vasculature, such as descending aorta and right carotid artery (RCA), of these mice (Figure 3B). Imaging of tissue sections confirmed these observations by showing that DiD signal was primarily delivered to the thickened intima of the LCA, with little-to-no fluorescence detected in the normal intima of the RCA (Figure 3C). These results provide strong evidence that Mo membrane cloaking enhances the interaction of the resulting particles with inflamed endothelium but not quiescent EC of the vasculature. Moreover, the biodistribution analysis showed that compared to the uncoated NP, MoNP-mediated delivery increased the fluorescent payload in the plasma and spleen and decreased its accumulation in the liver (Figure 3D). These findings indicated that while MoNP and NP have similar clearance routes through the liver and spleen, MoNP have extended residence time in circulation. Importantly, MoNP enables site-specific delivery of the payload to the inflamed endothelium displayed over plaque tissues, highlighting the potential of MoNP as a targeted delivery approach for atherosclerosis.

**Figure 3.**
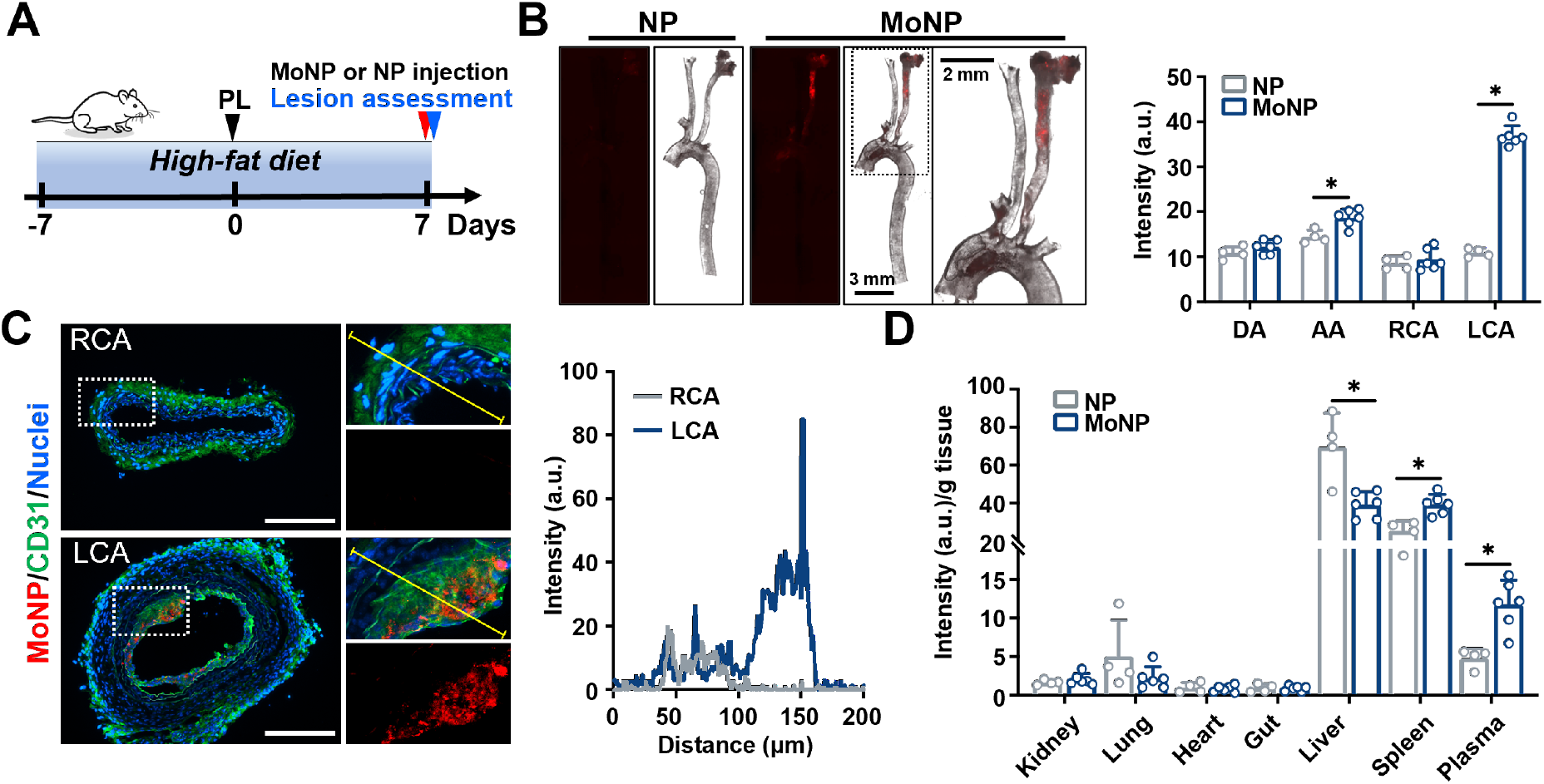
MoNP enabled active targeting of atherosclerotic vasculature. (A) A schematic showing the experimental design of MoNP-DiD or NP-DiD administration. (B) Representative fluorescent images of the arterial tissues isolated from ApoE^−/−^ mice receiving MoNP-DiD or NP-DiD, with quantification of the fluorescent intensity measured in LCA, RCA, aortic arch (AA), and descending aorta (DA). Red: MoNP-DiD or NP-DiD. (C) Representative fluorescent images of RCA and LCA cross-sections from the mouse receiving MoNP-DiD. Red: MoNP-DiD, green: CD31 and elastic layer autofluorescence, and blue: nuclei. The graph indicates the intensity of DiD signal of the crosslines (yellow). Scale bar = 200 μm. (D) The fluorescent intensity of the major organs isolated from ApoE^−/−^ mice receiving MoNP-DiD or NP-DiD. n = 6 for MoNP-DiD, and n = 4 for NP-DiD. *p < 0.05 vs. the NP-DiD group.

### 2.4. MoNP-VP treatment alleviates the TNFα-induced inflammatory responses in EC

Having established MoNP as an effective targeted delivery vehicle, we proceeded to investigate its potential in formulating a nanodrug aimed at blocking YAP/TAZ dysregulation in activated endothelium and assess its efficacy in reducing arterial inflammation and plaque development. To this end, we prepared MoNP loaded with VP (i.e., MoNP-VP) to suppress YAP/TAZ-TEAD interactions and tested the formulation in EC (Figure 4A). MoNP-VP has a hydrodynamic size of 289.4 nm, and approximately 75% of the input VP was loaded into MoNP, with a LE of 16.8% (Figure 4B). EC treated with MoNP-VP exhibited significantly decreased YAP/TAZ expression and inflammatory responses induced by TNFα, compared to the vehicle control (i.e., MoNP), as evidenced by reductions of CTGF, VCAM1, and ICAM1 expressions and Mo attachment to EC (Figure 4C, 4D, and 4E). Moreover, we performed RNA-sequencing (RNA-seq) profiling to assess the transcriptomic responses of EC to MoNP-VP treatment compared to MoNP, which identified a total of 4,707 differentially expressed genes (DEGs), with 2,333 upregulated and 2,374 downregulated (Figure S7A). Consistently, MoNP-VP reduced the expressions of atheroprone genes, including VCAM1, ICAM1, SELE, CCL2, and CXCR4, as well as YAP/TAZ-targeted genes such as CTGF, GATA3, and TNS3 (Figure 4F and S7B). Additionally, MoNP-VP increased the expressions of atheroprotective genes, including KLF2, KLF4, and NRF2. We further performed the Kyoto Encyclopedia of Genes and Genomes (KEGG) pathway enrichment and Gene Ontology (GO) analysis using ClusterProfiler (Figure 4G, S7C, and S7D). Our results identified TNF signaling pathway, NF-κB signaling pathway, cytokine-cytokine receptor interaction, fluid shear stress and atherosclerosis, and Hippo signaling pathway among the top enriched pathways, indicating MoNP-VP treatment exerted significant effects on the molecular pathways involved in endothelial inflammation and atherosclerosis. These results, collectively, suggest that MoNP-VP can effectively suppress YAP/TAZ activation and subsequent atherogenic responses in EC.

**Figure 4.**
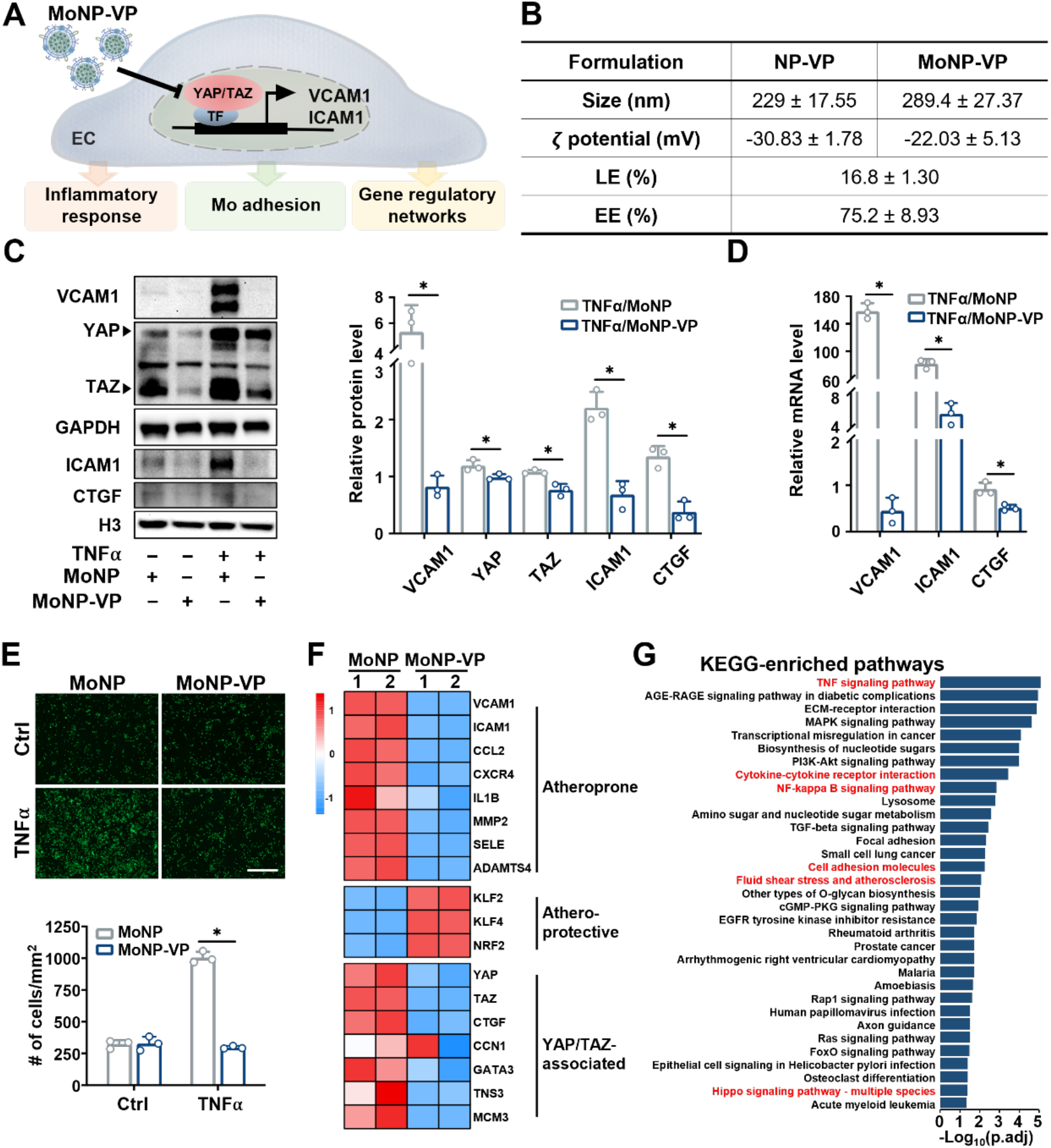
MoNP-VP treatment alleviated the inflammatory response in EC. (A) A schematic showing the experimental design of MoNP-VP treatment in EC. (B) Physicochemical characterization of MoNP-VP. (C) Western blot analysis of the TNFα-induced expression of VCAM1, ICAM1, YAP/TAZ, and CTGF in EC treated with MoNP-VP or MoNP. (D) qRT-PCR analysis of the TNFα-induced expression of VCAM1, ICAM1, and CTGF in EC treated with MoNP-VP or MoNP. For C and D, the data are normalized to its respective loading controls and the MoNP group. n = 3; *p < 0.05 vs. TNFα/MoNP. (E) Representative images of fluorescently labeled Mo (green) attached to EC monolayers treated with MoNP-VP or MoNP. Scale bar = 500 μm. N = 3; *p < 0.05 vs. TNFα/MoNP. (F) The RNA-Seq heatmap results displayed differences in the expression of atheroprone, atheroprotective, and YAP/TAZ-associated genes in TNFα-stimulated EC pretreated with MoNP-VP compared to those with MoNP. The genes are ranked based on their z-scores. (G) The KEGG pathway enrichment analysis of DEGs in response to MoNP-VP vs. MoNP in TNFα-stimulated EC. The inflammatory-related pathways and Hippo signaling pathway were highlighted in red. padj. < 0.05.

### 2.5. MoNP-VP treatment attenuates arterial inflammation and atherosclerosis *in vivo*

We next evaluated the pharmacological actions of MoNP-VP in mouse arteries. Specifically, we assessed the impact of MoNP-VP treatment on YAP/TAZ expression and arterial inflammation using the PL model. ApoE^-/-^ mice were intravenously administered with MoNP-VP or MoNP at a dose of 2 mg/kg immediately after PL and then every 72 hours for a total of 3 injections (Figure 5A). We collected the arterial tissue 1 week after the operation to assess the effects of MoNP-VP on YAP/TAZ expression and macrophage recruitment. Consistent with our *in vitro* data, the expressions of YAP/TAZ and CTGF in the arterial wall were significantly decreased by MoNP-VP treatment, as compared to MoNP (Figure 5B). Furthermore, immunofluorescence staining of LCA cross-sections demonstrated that MoNP-VP treatment markedly reduced YAP/TAZ expression throughout the vessel wall, VCAM1 expression in the intima layer, and the number of infiltrated CD68-positive cells, compared to MoNP (Figure 5C). These findings indicate that VP delivered by MoNP is highly effective in blocking YAP/TAZ expression in vascular cells, subsequently reducing PL-induced arterial inflammation by suppression of endothelial activation and leukocyte recruitment.

**Figure 5.**
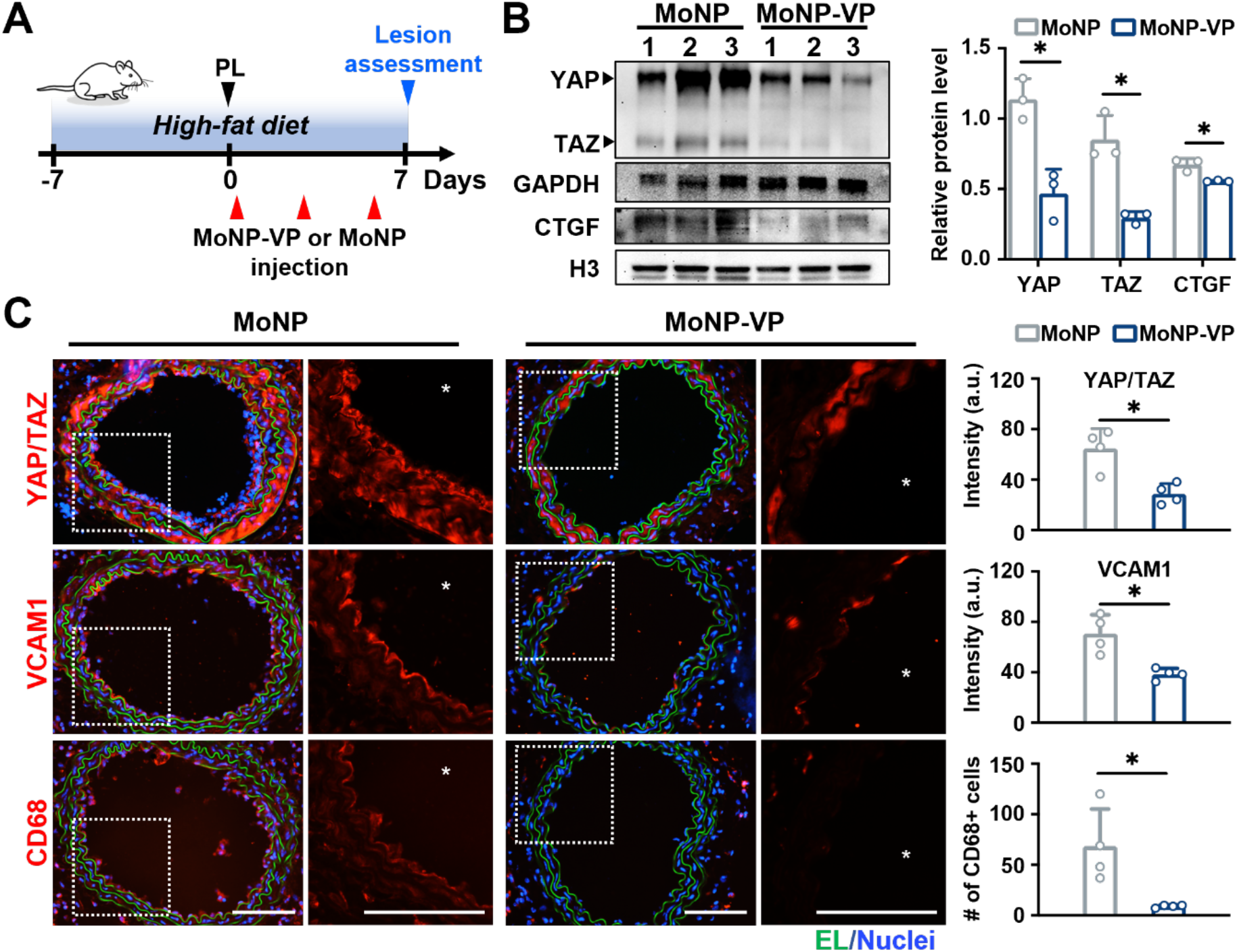
MoNP-VP treatment suppressed the PL-induced inflammatory response *in vivo*. (A) A schematic showing the experimental design of MoNP-VP treatment (2 mg/kg) in ApoE^−/−^ mice. The arterial tissues were harvested 7 days after the PL procedure. (B) Western blot analysis of the expressions of YAP/TAZ and CTGF in response to MoNP-VP or MoNP treatment. The band intensity is normalized to its respective loading controls. n = 3; *p < 0.05 vs. MoNP (C) Immunofluorescence staining of YAP/TAZ, VCAM1, and CD68-positive cells in the arterial wall. Red: YAP/TAZ, VCAM1, or CD68, green: elastic layer autofluorescence, blue: nuclei. White asterisks indicate the lumen. The intensity of YAP/TAZ and VCAM1 in the intimal layer of LCA and the number of infiltrated CD68-positive cells were quantified. n = 4, *p < 0.05 vs. MoNP.

To assess the therapeutic potential of MoNP-VP treatment in plaque development, we administered MoNP-VP, MoNP, free VP, or saline to ApoE^-/-^ mice that underwent PL for a total of 6 injections. Four weeks after the PL procedure, we harvested the arterial tissue and serum to analyze the size of plaque buildup in the LCA and total cholesterol level, respectively (Figure 6A). Only MoNP-VP treatment, but not free VP or MoNP, was able to reduce the overall size of atherosclerotic lesions in the LCA (Figure 6B). To quantify plaque size, we analyzed three segments of the LCA from the proximal end to the carotid bifurcation using Oil Red O and hematoxylin staining of LCA cross-sections. Compared to the free VP treatment and controls (i.e., MoNP, saline), MoNP-VP treatment significantly decreased PL-induced carotid atherosclerosis, as evidenced by the reductions of Oil Red O-positive plaque area and percentage of luminal occlusion in the LCA (Figure 6C, 6D, 6E, and S8). These results strongly support that MoNP-VP, without photoactivation, effectively suppressed plaque development at a dose of 2 mg/kg and that our targeted delivery strategy significantly improved outcomes compared to the systemic administration of VP, at least within the tested dosing regimen. Additionally, there was no difference in the level of serum cholesterol between MoNP-VP and other groups, suggesting MoNP-VP exerts direct anti-atherosclerotic effects on the affected artery, rather than modulation of blood cholesterol (Figure 6F).

**Figure 6.**
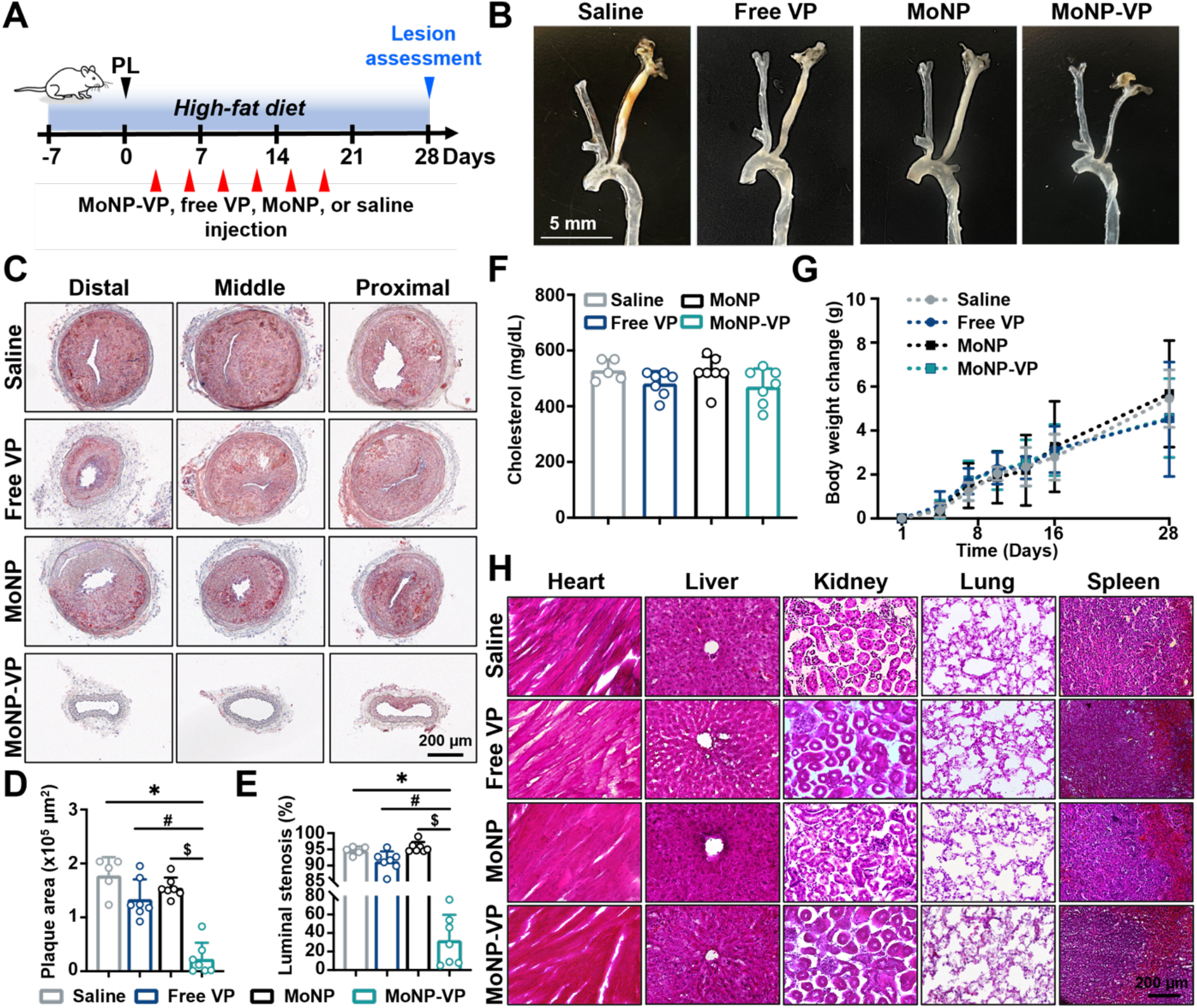
MoNP-VP treatment attenuated plaque development in ApoE^−/−^ mice. (A) A schematic showing the long-term treatment of MoNP-VP in mouse carotid atherosclerosis. ApoE^−/−^ mice were subjected to the PL procedure, followed by intravenous administration of MoNP-VP, free VP, MoNP, or saline after PL, and every 72 hours afterward, for a total of 6 injections. The lesion in the LCA was assessed on day 28 after PL. (B) Representative images of the arterial tissues (from carotid bifurcation to DA) of various treatment groups. (C) Representative images of the distal, middle, and proximal segments of the partially ligated LCA cross-sections from various treatment groups stained with Oil Red O and hematoxylin. (D) The Oil Red O-positive area and (E) the degree of luminal stenosis of the LCA were quantified, respectively. (F) The level of total cholesterol in mouse serum samples was measured. (G) The changes in body weight of the mice throughout the experiment. (H) Histological analysis of major organs isolated from ApoE^−/−^ mice subjected to various treatments. n = 7 each for MoNP-VP, free VP, and MoNP, and n = 5 for saline. *p < 0.05 vs. saline, ^#^p < 0.05 vs. free VP, and ^$^p < 0.05 vs. MoNP.

As the assessment of potential toxicity, we monitored body weight of the mice throughout the experiment and did not observe significant change due to MoNP-VP treatment (Figure 6G). We also harvested major organs at the end of the experiment and did not detect obvious histopathological alterations in any of the organs at the tested dose and frequency. (Figure 6H). Additionally, we analyzed the serum samples using a metabolic panel test, which showed that MoNP-VP treatment did not alter levels of blood chemicals and metabolites, including markers indicating liver or kidney damage (Table S1). These findings demonstrated that the MoNP-VP dosage used in this experiment did not elicit apparent side effects in major organs.

## 3. Discussion

A promising strategy for treating atherosclerosis is locally blocking inflammatory pathways and cellular events in the arterial wall, potentially minimizing systemic exposure and unwanted side effects compared to systemic routes. Despite its promise, the development of localized atherosclerosis therapies has been hindered by the lack of a plaque-specific delivery vehicle and effective molecular targets. To address this unmet need, we have developed a novel lesion-targeted biomimetic nanodrug, MoNP, packaged with a therapeutic agent, VP, to suppress YAP/TAZ activities. Our *in vitro* and *in vivo* findings have validated that MoNP enhances the uptake of the payload by inflamed EC while avoiding phagocytic cells, thereby extending their circulation time and achieving lesion-specific delivery. As the proof-of-concept investigation, our study provides compelling evidence demonstrating that MoNP-mediated delivery of VP effectively reduces YAP/TAZ-mediated inflammatory events in EC and attenuates macrophage infiltration and plaque development in the arteries of ApoE^-/-^ mice without causing noticeable adverse reactions in major organs. These findings suggest that local inhibition of YAP/TAZ activities using MoNP-VP could represent a highly effective and safe strategy for treating arterial inflammation and atherosclerosis.

Despite being a critical pathological feature in atherosclerosis, the inflamed, activated EC offer a valuable avenue for targeted drug delivery to plaque sites. Indeed, several delivery vehicles leveraging peptides or antibodies targeting specific adhesion molecules on the surface of inflamed EC have been developed to guide the delivery of payloads to plaque sites [26–28]. However, challenges, such as the immunogenicity and stability of the vehicles, as well as the heterogeneity of adhesion molecule expression patterns in EC from different vascular regions, could impact the effectiveness of ligand-mediated delivery approaches [29,30]. Unlike traditional NP functionalization that may require complex conjugation processes, our biomimetic platform offers an innovative, one-step strategy to coat NP with a lipid bilayer membrane derived from primary Mo, which improves immune evasion and most importantly, enhances targeting capabilities. The resulting MoNP display multiple targeting moieties of classical Mo including integrins (e.g., very late antigen-4), CCR2, and selectins on their surface, which simultaneously interact with the adhesion molecules (e.g., VCAM1, ICAM1), ligands, and receptor on activated EC to achieve strong adherence. This is supported by our *in vitro* study showing that Mo membrane cloaking significantly enhanced the internalization of NP into TNFα-stimulated EC compared to untreated EC. The enhanced uptake of MoNP by activated EC could be blocked by pretreatment with a VCAM1 antibody, suggesting similar to the homing of circulating Mo, the inflammatory tropism of EC plays a primary role in recruiting MoNP to the diseased artery, alleviating concerns about unintended interactions with healthy vasculature. Furthermore, the selective targeting capability of MoNP to inflamed vasculature was further validated using the PL model, in which the fluorescent payload of MoNP was found to exclusively accumulate in the early plaque regions of LCA, highlighting its strong potential as an atherosclerosis-targeted drug delivery vehicle. Further exploration of the stability of MoNP in the arterial wall and their ability to transverse through the endothelium beyond initial binding would undoubtedly advance our understanding and application of this promising delivery strategy.

One major advantage of MoNP over other EC-targeted delivery vehicles is their potential to reduce the opsonization of nanocarriers by the MPS and extend the retention time of payloads in circulation. Our *in vitro* and *in vivo* results provide clear evidence for the enhanced immune evasion of MoNP, demonstrating that a significantly greater internalization of uncoated NP by phagocytes compared to MoNP after incubation, and that the plasma of mice receiving MoNP had a considerably higher concentration of fluorescent payload 4 hour-post administration. Despite these promising findings, further optimization may be needed to reduce liver and splenic filtration/removal of MoNP. Potential modifications of MoNP include increasing the membrane-to-NP ratio, improving membrane coating efficiency, and incorporating “don’t eat me” signals (e.g., CD47) or the membrane from myeloid cells with a longer half-life might be viable solutions [31]. Additionally, when translating MoNP for clinical application, using Mo membrane sourced from the patient’s own blood or bone marrow would minimize immunogenicity and extend blood resident time, ultimately enhancing therapeutic efficacy.

A critical challenge in treating plaque development and progression lies in the lack of druggable targets linked to inflammatory-fibrotic activities in the arterial wall. Building on the pro-atherogenic roles of YAP/TAZ dysregulation, we developed a novel therapeutic strategy employing MoNP for targeted delivery of a YAP/TAZ antagonist, which specifically confines YAP/TAZ inhibition to the artery affected by atherosclerosis. Recognizing its ability to disrupt YAP/TAZ-TEAD interactions and status as an FDA-approved agent, we chose non-photoactivated VP as a therapeutic payload to treat YAP/TAZ-induced pathologies in atherosclerotic arteries. In line with the literature demonstrating VP’s potential in cancer therapy, we found that MoNP-VP treatment induced YAP/TAZ degradation in cultured EC in a light-independent manner, leading to the suppression of TNFα-induced inflammatory phenotypes [7,15]. Our RNA-seq assay further supports that MoNP treatment effectively suppressed inflammatory responses in EC through dual effects: downregulating YAP/TAZ-associated and atheroprone genes, and upregulating atheroprotective genes. Additionally, our KEGG pathway enrichment analysis of the RNA-seq data reveals that VP delivered by MoNP exerts inhibitory effects primarily on inflammatory pathways associated with atherosclerosis, including Hippo-YAP/TAZ signaling, without affecting those involved in cell necrosis or apoptosis. Collectively, these findings establish MoNP-VP as a lesion-targeted, pathway-specific nanodrug that effectively blocks YAP/TAZ functions and mitigates inflammatory responses in EC while preserving essential functions. This underscores the potential for repurposing VP as a therapeutic option in the treatment of atherosclerosis and other inflammatory diseases.

Emerging evidence has demonstrated the potential of VP for photodynamic therapy in the treatment of cancers [16]. In the context of atherosclerosis, Jain et al. reported that upon photoactivation, VP produced reactive oxygen species in atherosclerotic arteries isolated from mice, which induced apoptosis of immune and vascular cells in the arterial wall and decreased vasocontraction [32]. However, the effects of non-photoactivated VP on atherosclerosis remain completely unknown. This present study provides the first preclinical evidence highlighting the therapeutic potential of our targeted VP nanodrug in treating arterial inflammation and atherosclerosis through a light-independent mechanism. Specifically, our animal studies showed that without photoactivation, MoNP-VP treatment at a dose of 2 mg/kg was effective to reduce the expression of YAP/TAZ and VCAM1 in PL-operated arteries and macrophage infiltration. In a long-term experiment, repeated doses of MoNP-VP greatly diminished PL-induced plaque formation without altering the serum lipid profile, whereas the free drug counterpart had little-to-no effect on plaque sizes. This suggests that targeted delivery is more efficient than systemic routes, as a much higher dose of VP may be required for the latter to achieve a therapeutic effect. Although the present study focused primarily on the impact of MoNP-VP on inflammatory responses in EC and plaque development, it is worth noting that YAP/TAZ activation and overexpression have also been associated with the dedifferentiation of vascular smooth muscle cells (SMC), as well as the polarization of macrophages towards a proinflammatory M1 phenotype, known to exacerbate plaque progression [33–36]. In this regard, the locally released VP in the diseased arterial wall could potentially exert additional beneficial effects on suppressing the pathological phenotypes of SMC and reducing the number of M1 macrophages, further improving the treatment outcome for atherosclerosis. One limitation of our study is that the anti-atherosclerotic effects of MoNP-VP were examined on a model of plaque development. Future investigation is warranted to evaluate the therapeutic efficacy of MoNP-VP in ameliorating pre-existing or advanced lesions, as well as to unravel the mechanisms underlying pharmacological effects on vascular and immune cells in plaque stabilization or regression.

A successful therapeutic must demonstrate both efficacy and safety. Although the dosage of VP we used was significantly lower than those reported in the literature, and MoNP-mediated delivery could minimize systemic exposure, there is still potential organ toxicity induced by VP [17,37,38]. To assess this, we conducted histology staining and serum biochemistry analyses, which revealed no significant pathological characteristics in major organs attributable to MoNP-VP treatment. Moreover, the body weight changes observed in all experimental mice remained within the normal range. These findings suggest that the dose utilized in our study is well-tolerated in mice, and that the MoNP-VP treatment of atherosclerosis is both safe and therapeutically effective in alleviating arterial inflammation and plaque development. Nevertheless, further safety studies are necessary to assess potential chronic toxicity after long-term use in large animal models before proceeding to clinical trials.

## 4. Conclusion

In summary, we have successfully developed a novel atherosclerosis-targeted drug delivery system utilizing nanocarriers coated with Mo membrane. This innovative MoNP platform greatly enhances the immune evasion of the nanocarriers and boosts their targeting to inflamed vasculature while avoiding healthy blood vessels. We employed MoNP to deliver VP, achieving lesion-targeted and pathway-specific suppression of YAP/TAZ, leading to robust anti-inflammatory and anti-atherosclerotic effects without requiring photoactivation. Furthermore, MoNP-mediated delivery maximizes the therapeutic efficacy of VP while minimizing its organ toxicity. Our study paves the way to harness biomimicry and nanomedicine for more effective and tailored treatment options for atherosclerosis and other inflammatory diseases, propelling the advancement of personalized and precision medicine.

## 5. Experimental Section

### Isolation of bone marrow-derived Mo and Mo vesicles

Mouse Mo were derived from BMC isolated from the femurs, humerus, and tibiae of C57BL/6 mice as described [39]. Briefly, BMC were flushed out from the bone and cultured in RPMI 1640 medium with 10% FBS, 1% penicillin-streptomycin, 1% L-glutamate, 1% sodium pyruvate, and 20 ng/mL M-CSF for 5 days. Mo were then isolated from the suspension of differentiated BMC using a MojoSort™ Mouse Monocyte Isolation Kit (BioLegend #480154). The expression of Mo surface markers was measured by flow cytometric analysis using fluorescently labeled antibodies against CD11b (BioLegend #101207), CD45R (BioLegend #103211), Ly6C (BioLegend #128021), and CCR2 (BioLegend #150627). To obtain Mo membrane vesicles, the plasma membrane fraction was separated using a Minute™ Plasma Membrane Protein Isolation and Cell Fractionation kit (Invent Biotechnologies).

### Preparation and characterization of MoNP

NP were prepared using an emulsion-solvent evaporation method described previously [22]. Briefly, 10 mg of poly(D, L-lactide-co-glycolide) (PLGA; Resomer^®^ RG 503H, Sigma Aldrich) was dissolved in 1 mL of dichloromethane (DCM, Sigma Aldrich) and added dropwise to 2% polyvinyl-alcohol (PVA, Acros Organics) solution. The resulting emulsion was sonicated with the probe sonicator (Fisherbrand) and added to a 0.5% PVA solution, then stirred for 3 hours to remove the solvent. NP were washed and collected by centrifugation. For the encapsulation of payloads, 400 μg of DiD (Biotium) or 2 mg of VP (Tocris Bioscience) was dissolved in 1 mL of DCM containing PLGA. To prepare MoNP, the isolated Mo vesicles were added with NP at a membrane protein-to-NP weight ratio of 1:10 and mixed in an ultrasonic bath for 5 minutes. The hydrodynamic size, PDI, and zeta potential of resulting particles were measured using DLS Zetasizer (Malvern). The morphology of MoNP and NP was examined after staining with 0.2% uranyl acetate using the TEM (Philips CM12) at ASU Eyring Materials Center. To determine the encapsulation efficiency (EE) and LE, NP loaded with DiD or VP were lyophilized and then dissolved in dimethylsulfoxide (DMSO), followed by the measurement of fluorescent intensity using a plate reader (BioTek). EE and LE were calculated as follows:

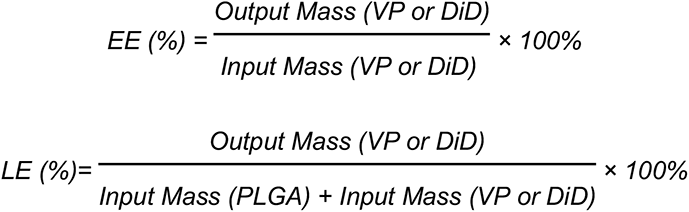

### *In vitro* release of NP-DiD

The release of NP-DiD was assessed using the direct release method. Briefly, aliquots of NP-DiD were resuspended in three different solutions (saline pH 7, saline pH 7 containing 10% serum, and saline pH 6) and agitated at 100 rpm at 37°C for predetermined time intervals ranging from 1 to 96 hours. Following incubation, the samples were centrifugated and dissolved in DMSO. The fluorescent intensity of the samples was then measured using a plate reader. The cumulative percentage of DiD released was calculated using the equation: Cumulative dye release (%) = (Read(0) - Read(t))/Read(0) × 100), where Read(0) and Read(t) represent the amount of DiD loaded and the amount of DiD released at time t, respectively.

### Cellular uptake and intracellular fate of MoNP

Human umbilical cord vein EC (ATCC), Mo, and macrophages derived from mouse BMC were maintained in their respective growth medium under standard cell culture conditions (37°C, 5% CO_2_, 100% humidity). For the cellular uptake assays, MoNP-DiD or NP-DiD were incubated with untreated EC, TNFα-pretreated EC, TNFα-/anti-VCAM1-pretreated EC, Mo, and macrophages for 30 minutes and then replaced with fresh medium. For the SS experiment, MoNP-DiD were added to the circulating medium in a parallel plate flow system and allowed to interact with EC exposed to high or low SS (12 or 1 dyne/cm^2^) for 2 hours [40]. The intracellular DiD signal was quantified using fluorescent imaging and flow cytometric analyses. For the colocalization study, MoNP- or NP-treated EC were stained with LysoTracker (Cell Signaling) and DAPI and observed using the confocal microscope (Leica SP8) at ASU Advanced Light Microscopy Facilities. Colocalization of MoNP-DiD or NP-DiD with lysosomes was determined using the JACoP plugin in ImageJ on randomly selected cells and calculation of Pearson’s correlation coefficients [41].

### Biocompatibility testing

To assess cytotoxicity of MoNP, EC were incubated with MoNP, NP, or PBS. The number of live EC was quantified on day 1, 2, and 3 post-seeding. Hemolysis induced by MoNP or NP was evaluated *in vitro* using the direct contact method. Briefly, mouse blood was diluted with saline and mixed with MoNP, NP, or deionized water as a control. The samples were incubated at 37°C for 1 hour and visually inspected after low-speed centrifugation. The supernatant absorbance was measured using a plate reader at 540 nm. For *in vivo* toxicity, ApoE^-/-^ mice were intravenously administered with MoNP or saline every 72 hours for a total of three injections. On day 9, the organs were harvested, sectioned, and stained with hematoxylin and eosin for histopathological analysis.

### SDS-PAGE and Western blot analysis

Equal amounts of protein were separated by sodium dodecyl sulfate-polyacrylamide gel electrophoresis (SDS-PAGE) and stained with a Gel-FAST Gel Staining/Destaining kit (BioVision) for visualization. For Western blot analysis, the proteins were transferred to nitrocellulose membranes after electrophoresis, followed by blocking with 5% BSA and incubation with primary and secondary antibodies. The following antibodies were used in this study at a dilution of 1:1000: CD11b (Cell Signaling #17800), Na^+^/K^+^ ATPase (Cell Signaling #3010), VCAM1 (Cell Signaling #13622), ICAM1 (Cell Signaling #4915), and GAPDH (Cell Signaling #2118), TLR4 (BioLegend #145401), Histone 3 (BioLegend #819411), YAP/TAZ (Santa Cruz #sc-101199), and CTGF (Novus Biologicals #NB100-724SS). The blots were developed using a chemiluminescence kit (Pierce) and imaged using the Analytik Jena bioimaging system. The band intensity was quantified using ImageJ. Uncropped Western blot images are displayed in Figure S9.

### RNA extraction and analysis

Total RNA was extracted using TRIzol reagent (Invitrogen) according to the manufacturer’s protocols. Subsequently, mRNA was reverse-transcribed to cDNA by using oligo-dT primer (Promega) and M-MLV reverse transcriptase (Promega). Quantitative real-time PCR (qRT-PCR) was performed on a QuantStudio 3 Real-Time PCR System (Applied Biosystems). Relative gene expression was determined by the 2^-ΔΔCT^ method, and the primer sets used for qRT-PCR are listed in Table S2. For RNA-seq transcriptome profiling, the RNA library and transcriptome sequencing was conducted by Novogene. The sequence read pairs were aligned to the human genome (hg38) using STAR, and the number of reads for each gene was counted using StringTie [42,43]. DESeq2 was used to identify DEGs, and genes with the sum of counts less than 100 across all samples were removed before the analysis [44]. The heatmap was generated by pheatmap (version 1.0.12) with a row z-score value. Genes with a multiple hypotheses-corrected P value of <0.05 and |log_2_FoldChange| ≥ 0.75 were considered DEGs. DEGs were subjected to ClusterProfiler, including the KEGG pathway and GO analysis [45–47]. The RNA-seq data are available in GEO DataSets with the accession number GSE229918.

### Mo adhesion assay

THP-1 cells were labeled with CellBrite® Cytoplasmic Membrane Dye (Biotium) and resuspended to the concentration of 5 × 10^5^ cells/mL. The labeled THP-1 cells were then added to TNFα-stimulated or control EC monolayers pretreated with MoNP-VP or MoNP. After incubation, unattached THP-1 cells were washed out with fresh medium. Fluorescent images showing Mo attachment to EC monolayers were taken under Lionheart FX Automated Microscope, and the number of adherent THP-1 cells was quantified using ImageJ.

### Animal experiments

All animal experiments performed in this study have been approved by the Institutional Animal Care and Use Committee (IACUC) of Arizona State University, USA. Male ApoE^-/-^ mice (8-12 weeks old), purchased from The Jackson Laboratory, were subjected to the PL procedure to induce inflammation and carotid atherosclerosis in the LCA [25]. All mice were fed a high-fat diet (Envigo #TD.88137) for 1 week prior to the PL procedure and continued to be fed the same diet until the time of euthanasia. The PL was conducted by ligating the left external CA, left internal CA, and left occipital artery while maintaining the left superior thyroid artery intact. For biodistribution analysis, MoNP-DiD was administered via retro-orbital injection to mice that underwent the PL 1 week prior. The arterial tissue (from carotid bifurcation to abdominal aorta) and organs were harvested 4 hours after the injection, and DiD intensity was quantified for each of the three individual zones within the arterial segments using fluorescent imaging. Subsequently, the left common CA was embedded in Tissue-Tek O.C.T. compound, cryosectioned, and stained with a CD31 antibody (R&D Systems #AF3628) and DAPI. For organ assessment, various tissues (kidney, lung, heart, gut, liver, and spleen) and plasma were homogenized, and the fluorescent intensity in homogenized tissues was measured. For the short-term experiment, MoNP-VP at a dose of 2 mg/kg was administered intravenously three times a week to assess the effect on PL-induced inflammatory response. Arterial tissues were harvested 24 hours after the 3rd injection of MoNP-VP for Western blot and immunofluorescence analysis using antibodies against YAP/TAZ, CTGF, VCAM1, and CD68 (BioLegend #137001). To evaluate the long-term effect on plaque development, the mice underwent PL and were then injected with MoNP-VP, free VP, MoNP, and saline via intravenous injection every 3 days for a total of 6 injections. Four weeks after PL, arterial tissues were harvested for lesion assessment. The LCA was cryosectioned and stained with Oil Red O and hematoxylin to evaluate the plaque size and luminal stenosis. Major organs were harvested, cryosectioned, and stained with hematoxylin and eosin for histopathological examination. Serum samples isolated from the mice were analyzed to determine their lipid profile and metabolic panel. These analyses were carried out by Protatek Reference Laboratory (Mesa, AZ).

### Statistical analysis

All graphical data are presented as mean ± standard deviation (SD), and statistical analysis from at least three biological repeat experiments were performed using GraphPad Prism 8.0. For comparisons between two groups, a two-tailed unpaired Student’s t-test was used. One-way ANOVA was performed for the comparison of multiple groups, and statistical significance among multiple groups was determined by Tukey’s multiple comparisons test.

## Supporting Information

Supporting Information is available from the Wiley Online Library.

## Data Availability statement

The data that support the findings of this study are available from the corresponding author upon reasonable request.

## Acknowledgments

H.-C.H. and T.-Y.W. contributed equally to this study. This work was supported by funding from National Institutes of Health grants HL135416 to K.-C.W. and HL145170 and HL108735 to Z.B.C. The authors thank Dr. Miyeko Mana and Clarissa Hoffman for their technical assistance.

## References

[1] P. Libby, P.M. Ridker, G.K. Hansson, Progress and challenges in translating the biology of atherosclerosis, Nature. 473 (2011) 317–325. https://doi.org/10.1038/nature10146.

[2] M. Bäck, A. Yurdagul, I. Tabas, K. Öörni, P.T. Kovanen, Inflammation and its resolution in atherosclerosis: mediators and therapeutic opportunities, Nat Rev Cardiol. 16 (2019) 389–406. https://doi.org/10.1038/s41569-019-0169-2.

[3] P.M. Ridker, B.M. Everett, T. Thuren, J.G. MacFadyen, W.H. Chang, C. Ballantyne, F. Fonseca, J. Nicolau, W. Koenig, S.D. Anker, J.J.P. Kastelein, J.H. Cornel, P. Pais, D. Pella, J. Genest, R. Cifkova, A. Lorenzatti, T. Forster, Z. Kobalava, L. Vida-Simiti, M. Flather, H. Shimokawa, H. Ogawa, M. Dellborg, P.R.F. Rossi, R.P.T. Troquay, P. Libby, R.J. Glynn, CANTOS Trial Group, Antiinflammatory Therapy with Canakinumab for Atherosclerotic Disease, N Engl J Med. 377 (2017) 1119–1131. https://doi.org/10.1056/NEJMoa1707914.

[4] S.M. Nidorf, A.T.L. Fiolet, A. Mosterd, J.W. Eikelboom, A. Schut, T.S.J. Opstal, S.H.K. The, X.-F. Xu, M.A. Ireland, T. Lenderink, D. Latchem, P. Hoogslag, A. Jerzewski, P. Nierop, A. Whelan, R. Hendriks, H. Swart, J. Schaap, A.F.M. Kuijper, M.W.J. van Hessen, P. Saklani, I. Tan, A.G. Thompson, A. Morton, C. Judkins, W.A. Bax, M. Dirksen, M. Alings, G.J. Hankey, C.A. Budgeon, J.G.P. Tijssen, J.H. Cornel, P.L. Thompson, LoDoCo2 Trial Investigators, Colchicine in Patients with Chronic Coronary Disease, N Engl J Med. 383 (2020) 1838–1847. https://doi.org/10.1056/NEJMoa2021372.

[5] R.A. Baylis, D. Gomez, Z. Mallat, G. Pasterkamp, G.K. Owens, The CANTOS Trial: One Important Step for Clinical Cardiology but a Giant Leap for Vascular Biology, Arterioscler Thromb Vasc Biol. 37 (2017) e174–e177. https://doi.org/10.1161/ATVBAHA.117.310097.

[6] Y. Niu, N. Bai, Y. Ma, P.-Y. Zhong, Y.-S. Shang, Z.-L. Wang, Safety and efficacy of anti-inflammatory therapy in patients with coronary artery disease: a systematic review and meta-analysis, BMC Cardiovasc Disord. 22 (2022) 84. https://doi.org/10.1186/s12872-022-02525-9.

[7] K.-C. Wang, Y.-T. Yeh, P. Nguyen, E. Limqueco, J. Lopez, S. Thorossian, K.-L. Guan, Y.-S.J. Li, S. Chien, Flow-dependent YAP/TAZ activities regulate endothelial phenotypes and atherosclerosis, Proc Natl Acad Sci U S A. 113 (2016) 11525–11530. https://doi.org/10.1073/pnas.1613121113.

[8] L. Wang, J.-Y. Luo, B. Li, X.Y. Tian, L.-J. Chen, Y. Huang, J. Liu, D. Deng, C.W. Lau, S. Wan, D. Ai, K.-L.K. Mak, K.K. Tong, K.M. Kwan, N. Wang, J.-J. Chiu, Y. Zhu, Y. Huang, Integrin-YAP/TAZ-JNK cascade mediates atheroprotective effect of unidirectional shear flow, Nature. 540 (2016) 579–582. https://doi.org/10.1038/nature20602.

[9] P. Yuan, Q. Hu, X. He, Y. Long, X. Song, F. Wu, Y. He, X. Zhou, Laminar flow inhibits the Hippo/YAP pathway via autophagy and SIRT1-mediated deacetylation against atherosclerosis, Cell Death Dis. 11 (2020) 141. https://doi.org/10.1038/s41419-020-2343-1.

[10] T. Panciera, L. Azzolin, M. Cordenonsi, S. Piccolo, Mechanobiology of YAP and TAZ in physiology and disease, Nat Rev Mol Cell Biol. 18 (2017) 758–770. https://doi.org/10.1038/nrm.2017.87.

[11] S. Wang, L. Zhou, L. Ling, X. Meng, F. Chu, S. Zhang, F. Zhou, The Crosstalk Between Hippo-YAP Pathway and Innate Immunity, Front Immunol. 11 (2020) 323. https://doi.org/10.3389/fimmu.2020.00323.

[12] H.-J. Choi, N.-E. Kim, B.M. Kim, M. Seo, J.H. Heo, TNF-α-Induced YAP/TAZ Activity Mediates Leukocyte-Endothelial Adhesion by Regulating VCAM1 Expression in Endothelial Cells, Int J Mol Sci. 19 (2018) 3428. https://doi.org/10.3390/ijms19113428.

[13] K.J. Messmer, S.R. Abel, Verteporfin for age-related macular degeneration, Ann Pharmacother. 35 (2001) 1593–1598. https://doi.org/10.1345/aph.10365.

[14] Y. Liu-Chittenden, B. Huang, J.S. Shim, Q. Chen, S.-J. Lee, R.A. Anders, J.O. Liu, D. Pan, Genetic and pharmacological disruption of the TEAD-YAP complex suppresses the oncogenic activity of YAP, Genes Dev. 26 (2012) 1300–1305. https://doi.org/10.1101/gad.192856.112.

[15] J. Feng, J. Gou, J. Jia, T. Yi, T. Cui, Z. Li, Verteporfin, a suppressor of YAP-TEAD complex, presents promising antitumor properties on ovarian cancer, Onco Targets Ther. 9 (2016) 5371–5381. https://doi.org/10.2147/OTT.S109979.

[16] C. Wei, X. Li, The Role of Photoactivated and Non-Photoactivated Verteporfin on Tumor, Front Pharmacol. 11 (2020) 557429. https://doi.org/10.3389/fphar.2020.557429.

[17] K. Vigneswaran, N.H. Boyd, S.-Y. Oh, S. Lallani, A. Boucher, S.G. Neill, J.J. Olson, R.D. Read, YAP/TAZ Transcriptional Coactivators Create Therapeutic Vulnerability to Verteporfin in EGFR-mutant Glioblastoma, Clin Cancer Res. 27 (2021) 1553–1569. https://doi.org/10.1158/1078-0432.CCR-20-0018.

[18] R.H. Fang, A.V. Kroll, W. Gao, L. Zhang, Cell Membrane Coating Nanotechnology, Adv Mater. 30 (2018) e1706759. https://doi.org/10.1002/adma.201706759.

[19] S. Zou, B. Wang, C. Wang, Q. Wang, L. Zhang, Cell membrane-coated nanoparticles: research advances, Nanomedicine (Lond). 15 (2020) 625–641. https://doi.org/10.2217/nnm-2019-0388.

[20] C.-M.J. Hu, L. Zhang, S. Aryal, C. Cheung, R.H. Fang, L. Zhang, Erythrocyte membrane-camouflaged polymeric nanoparticles as a biomimetic delivery platform, Proc Natl Acad Sci U S A. 108 (2011) 10980–10985. https://doi.org/10.1073/pnas.1106634108.

[21] R.H. Fang, W. Gao, L. Zhang, Targeting drugs to tumours using cell membrane-coated nanoparticles, Nat Rev Clin Oncol. 20 (2023) 33–48. https://doi.org/10.1038/s41571-022-00699-x.

[22] C.-M.J. Hu, R.H. Fang, K.-C. Wang, B.T. Luk, S. Thamphiwatana, D. Dehaini, P. Nguyen, P. Angsantikul, C.H. Wen, A.V. Kroll, C. Carpenter, M. Ramesh, V. Qu, S.H. Patel, J. Zhu, W. Shi, F.M. Hofman, T.C. Chen, W. Gao, K. Zhang, S. Chien, L. Zhang, Nanoparticle biointerfacing by platelet membrane cloaking, Nature. 526 (2015) 118–121. https://doi.org/10.1038/nature15373.

[23] Y. Wang, K. Zhang, T. Li, A. Maruf, X. Qin, L. Luo, Y. Zhong, J. Qiu, S. McGinty, G. Pontrelli, X. Liao, W. Wu, G. Wang, Macrophage membrane functionalized biomimetic nanoparticles for targeted anti-atherosclerosis applications, Theranostics. 11 (2021) 164–180. https://doi.org/10.7150/thno.47841.

[24] D. Han, F. Wang, Z. Qiao, B. Wang, Y. Zhang, Q. Jiang, M. Liu, Y. Zhuang, Q. An, Y. Bai, J. Shangguan, J. Zhang, G. Liang, D. Shen, Neutrophil membrane-camouflaged nanoparticles alleviate inflammation and promote angiogenesis in ischemic myocardial injury, Bioact Mater. 23 (2023) 369–382. https://doi.org/10.1016/j.bioactmat.2022.11.016.

[25] D. Nam, C.-W. Ni, A. Rezvan, J. Suo, K. Budzyn, A. Llanos, D. Harrison, D. Giddens, H. Jo, Partial carotid ligation is a model of acutely induced disturbed flow, leading to rapid endothelial dysfunction and atherosclerosis, Am J Physiol Heart Circ Physiol. 297 (2009) H1535–1543. https://doi.org/10.1152/ajpheart.00510.2009.

[26] A. Kheirolomoom, C.W. Kim, J.W. Seo, S. Kumar, D.J. Son, M.K.J. Gagnon, E.S. Ingham, K.W. Ferrara, H. Jo, Multifunctional Nanoparticles Facilitate Molecular Targeting and miRNA Delivery to Inhibit Atherosclerosis in ApoE(-/-) Mice, ACS Nano. 9 (2015) 8885– 8897. https://doi.org/10.1021/acsnano.5b02611.

[27] Z. Zhou, C.-F. Yeh, M. Mellas, M.-J. Oh, J. Zhu, J. Li, R.-T. Huang, D.L. Harrison, T.-P. Shentu, D. Wu, M. Lueckheide, L. Carver, E.J. Chung, L. Leon, K.-C. Yang, M.V. Tirrell, Y. Fang, Targeted polyelectrolyte complex micelles treat vascular complications in vivo, Proc Natl Acad Sci U S A. 118 (2021) e2114842118. https://doi.org/10.1073/pnas.2114842118.

[28] P. Li, L. Jin, L. Feng, Y. Wang, R. Yang, ICAM-1-carrying targeted nano contrast agent for evaluating inflammatory injury in rabbits with atherosclerosis, Sci Rep. 11 (2021) 16508. https://doi.org/10.1038/s41598-021-96042-y.

[29] Z. Zhao, A. Ukidve, J. Kim, S. Mitragotri, Targeting Strategies for Tissue-Specific Drug Delivery, Cell. 181 (2020) 151–167. https://doi.org/10.1016/j.cell.2020.02.001.

[30] H. Gunawardana, T. Romero, N. Yao, S. Heidt, A. Mulder, D.A. Elashoff, N.M. Valenzuela, Tissue-specific endothelial cell heterogeneity contributes to unequal inflammatory responses, Sci Rep. 11 (2021) 1949. https://doi.org/10.1038/s41598-020-80102-w.

[31] Y. Wu, S. Wan, S. Yang, H. Hu, C. Zhang, J. Lai, J. Zhou, W. Chen, X. Tang, J. Luo, X. Zhou, L. Yu, L. Wang, A. Wu, Q. Fan, J. Wu, Macrophage cell membrane-based nanoparticles: a new promising biomimetic platform for targeted delivery and treatment, J Nanobiotechnology. 20 (2022) 542. https://doi.org/10.1186/s12951-022-01746-6.

[32] M. Jain, M. Zellweger, A. Frobert, J. Valentin, H. van den Bergh, G. Wagnières, S. Cook, M.-N. Giraud, Intra-Arterial Drug and Light Delivery for Photodynamic Therapy Using Visudyne®: Implication for Atherosclerotic Plaque Treatment, Front Physiol. 7 (2016) 400. https://doi.org/10.3389/fphys.2016.00400.

[33] X. Wang, G. Hu, X. Gao, Y. Wang, W. Zhang, E.Y. Harmon, X. Zhi, Z. Xu, M.R. Lennartz, M. Barroso, M. Trebak, C. Chen, J. Zhou, The induction of yes-associated protein expression after arterial injury is crucial for smooth muscle phenotypic modulation and neointima formation, Arterioscler Thromb Vasc Biol. 32 (2012) 2662–2669. https://doi.org/10.1161/ATVBAHA.112.254730.

[34] X. Feng, P. Liu, X. Zhou, M.-T. Li, F.-L. Li, Z. Wang, Z. Meng, Y.-P. Sun, Y. Yu, Y. Xiong, H.-X. Yuan, K.-L. Guan, Thromboxane A2 Activates YAP/TAZ Protein to Induce Vascular Smooth Muscle Cell Proliferation and Migration, J Biol Chem. 291 (2016) 18947–18958. https://doi.org/10.1074/jbc.M116.739722.

[35] X. Zhou, W. Li, S. Wang, P. Zhang, Q. Wang, J. Xiao, C. Zhang, X. Zheng, X. Xu, S. Xue, L. Hui, H. Ji, B. Wei, H. Wang, YAP Aggravates Inflammatory Bowel Disease by Regulating M1/M2 Macrophage Polarization and Gut Microbial Homeostasis, Cell Rep. 27 (2019) 1176–1189.e5. https://doi.org/10.1016/j.celrep.2019.03.028.

[36] M.M. Mia, D.M. Cibi, S.A.B. Abdul Ghani, W. Song, N. Tee, S. Ghosh, J. Mao, E.N. Olson, M.K. Singh, YAP/TAZ deficiency reprograms macrophage phenotype and improves infarct healing and cardiac function after myocardial infarction, PLoS Biol. 18 (2020) e3000941. https://doi.org/10.1371/journal.pbio.3000941.

[37] J. Kim, J.G. Shamul, S.R. Shah, A. Shin, B.J. Lee, A. Quinones-Hinojosa, J.J. Green, Verteporfin-Loaded Poly(ethylene glycol)-Poly(beta-amino ester)-Poly(ethylene glycol) Triblock Micelles for Cancer Therapy, Biomacromolecules. 19 (2018) 3361–3370. https://doi.org/10.1021/acs.biomac.8b00640.

[38] S.R. Shah, J. Kim, P. Schiapparelli, C.A. Vazquez-Ramos, J.C. Martinez-Gutierrez, A. Ruiz-Valls, K. Inman, J.G. Shamul, J.J. Green, A. Quinones-Hinojosa, Verteporfin-Loaded Polymeric Microparticles for Intratumoral Treatment of Brain Cancer, Mol Pharm. 16 (2019) 1433–1443. https://doi.org/10.1021/acs.molpharmaceut.8b00959.

[39] A. Francke, J. Herold, S. Weinert, R.H. Strasser, R.C. Braun-Dullaeus, Generation of mature murine monocytes from heterogeneous bone marrow and description of their properties, J Histochem Cytochem. 59 (2011) 813–825. https://doi.org/10.1369/0022155411416007.

[40] M.J. Davis, S. Earley, Y.-S. Li, S. Chien, Vascular mechanotransduction, Physiol Rev. 103 (2023) 1247–1421. https://doi.org/10.1152/physrev.00053.2021.

[41] S. Bolte, F.P. Cordelières, A guided tour into subcellular colocalization analysis in light microscopy, J Microsc. 224 (2006) 213–232. https://doi.org/10.1111/j.1365-2818.2006.01706.x.

[42] A. Dobin, C.A. Davis, F. Schlesinger, J. Drenkow, C. Zaleski, S. Jha, P. Batut, M. Chaisson, T.R. Gingeras, STAR: ultrafast universal RNA-seq aligner, Bioinformatics. 29 (2013) 15–21. https://doi.org/10.1093/bioinformatics/bts635.

[43] M. Pertea, G.M. Pertea, C.M. Antonescu, T.-C. Chang, J.T. Mendell, S.L. Salzberg, StringTie enables improved reconstruction of a transcriptome from RNA-seq reads, Nat Biotechnol. 33 (2015) 290–295. https://doi.org/10.1038/nbt.3122.

[44] M.I. Love, W. Huber, S. Anders, Moderated estimation of fold change and dispersion for RNA-seq data with DESeq2, Genome Biol. 15 (2014) 550. https://doi.org/10.1186/s13059-014-0550-8.

[45] T. Wu, E. Hu, S. Xu, M. Chen, P. Guo, Z. Dai, T. Feng, L. Zhou, W. Tang, L. Zhan, X. Fu, S. Liu, X. Bo, G. Yu, clusterProfiler 4.0: A universal enrichment tool for interpreting omics data, Innovation (Camb). 2 (2021) 100141. https://doi.org/10.1016/j.xinn.2021.100141.

[46] M. Kanehisa, M. Furumichi, M. Tanabe, Y. Sato, K. Morishima, KEGG: new perspectives on genomes, pathways, diseases and drugs, Nucleic Acids Res. 45 (2017) D353–D361. https://doi.org/10.1093/nar/gkw1092.

[47] M. Ashburner, C.A. Ball, J.A. Blake, D. Botstein, H. Butler, J.M. Cherry, A.P. Davis, K. Dolinski, S.S. Dwight, J.T. Eppig, M.A. Harris, D.P. Hill, L. Issel-Tarver, A. Kasarskis, S. Lewis, J.C. Matese, J.E. Richardson, M. Ringwald, G.M. Rubin, G. Sherlock, Gene ontology: tool for the unification of biology. The Gene Ontology Consortium, Nat Genet. 25 (2000) 25–29. https://doi.org/10.1038/75556.

