## Supporting information for "Lesion-specific suppression of YAP/TAZ by biomimetic nanodrug ameliorates atherosclerosis development"

**Title**

T.-Y. Wang

School for Engineering of Matter, Transport, and Energy

Arizona State University

Tempe, AZ 85287, USA

Z.B. Chen

Department of Diabetes Complications and Metabolism

Arthur Riggs Diabetes and Metabolism Research Institute

Beckman Research Institute, City of Hope

Duarte, CA 91010, USA

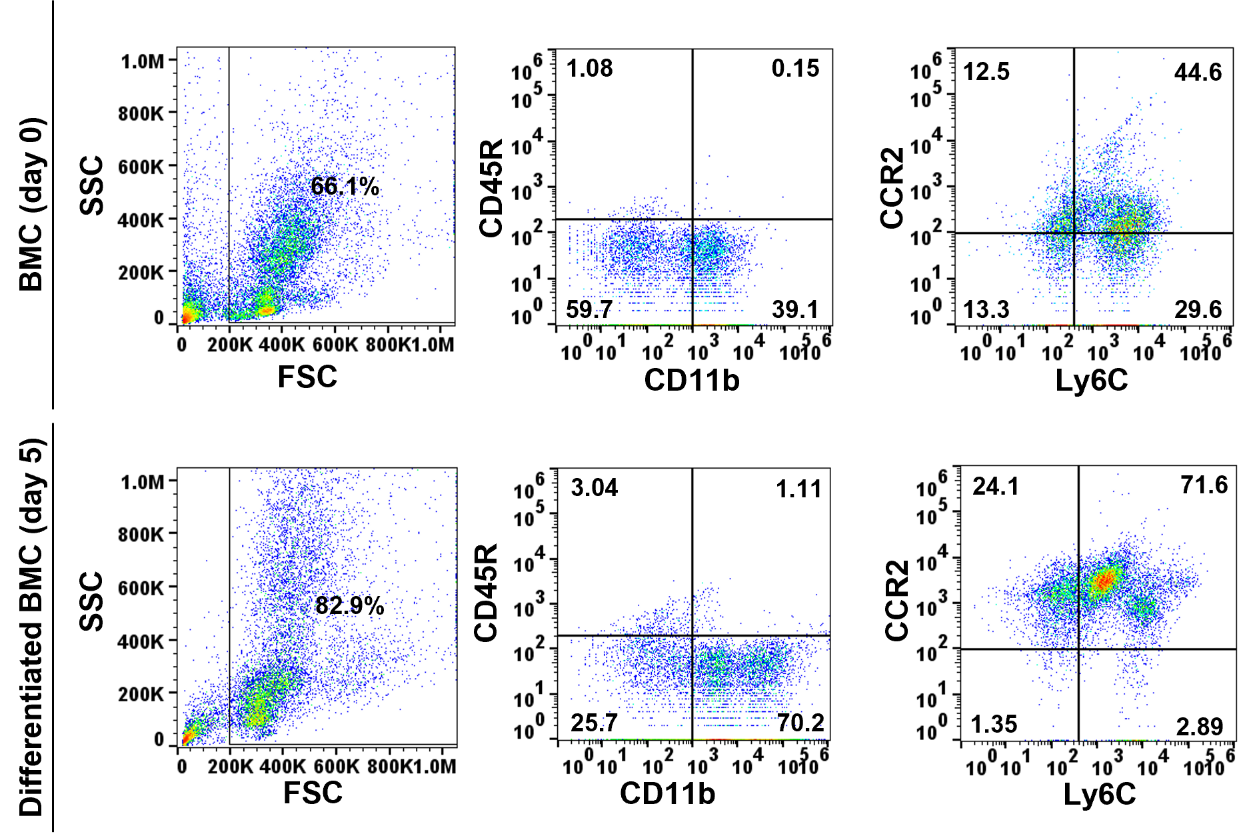

**Figure S1. Characterization of bone marrow cells (BMC) and M-CSF-differentiated BMC.** The expression of classical monocyte (Mo) markers (CD11b, Ly6C, and CCR2) in freshly isolated BMC and M-CSF-differentiated BMC was analyzed by flow cytometric analysis. The results are representative data from three independent assays.

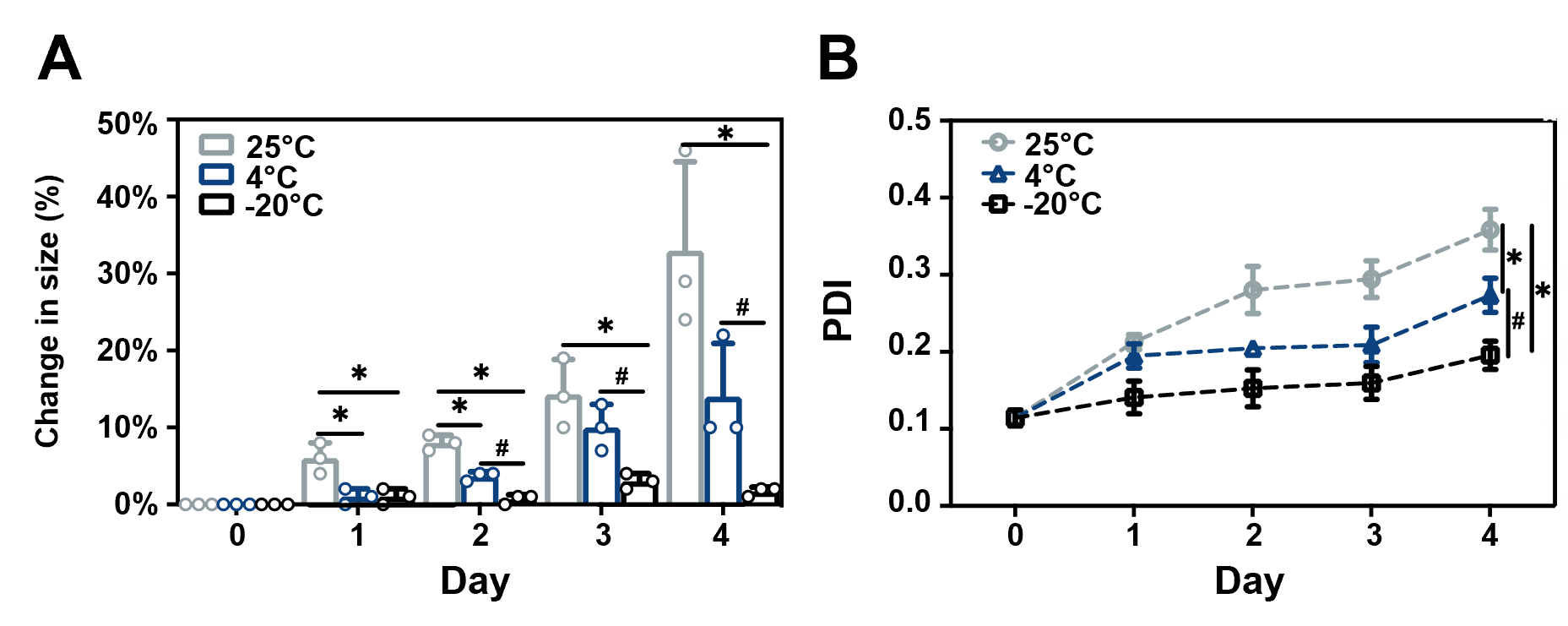

**Figure S2. *In vitro* stability of MoNP at different storage temperatures.** Mo membrane-coated nanoparticles (MoNP) were resuspended in the storage buffer (10% sucrose in saline) and incubated at 25°C, 4°C, or -20°C. The change in hydrodynamic size (A) and polydispersity index (PDI) (B) of MoNP were measured on day 1, 2, 3, and 4 after incubation using DLS analysis. n = 3. *p < 0.05 vs. 25°C and ^#^p < 0.05 vs. 4°C.

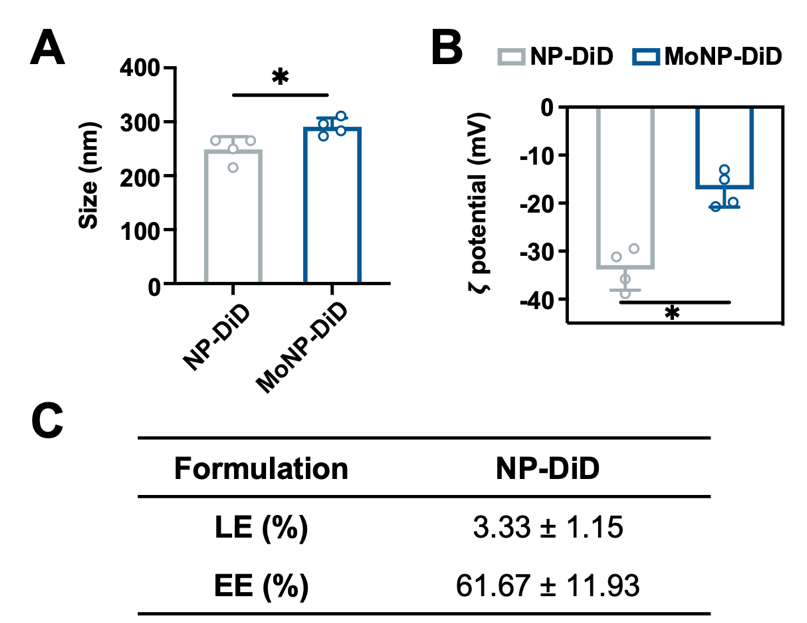

**Figure S3. Physicochemical characterization of MoNP-DiD.** The DLS results of MoNP-DiD and NP-DiD: (A) hydrodynamic size and (B) surface charge. (C) Loading efficiency (LE) and encapsulation efficiency (EE) of MoNP-DiD. n = 3-4, *p < 0.05 vs. NP-DiD.

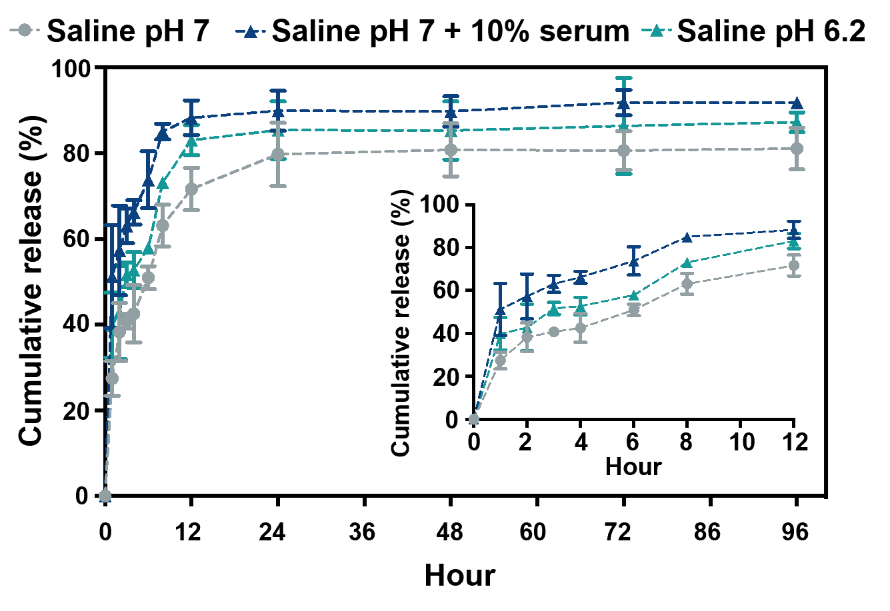

**Figure S4. *In vitro* payload release profiles of NP-DiD.** The cumulative release profile of NP-DiD in saline pH 7, saline pH 7 containing 10% serum, or saline pH 6 for 96 hours was assessed. n = 3.

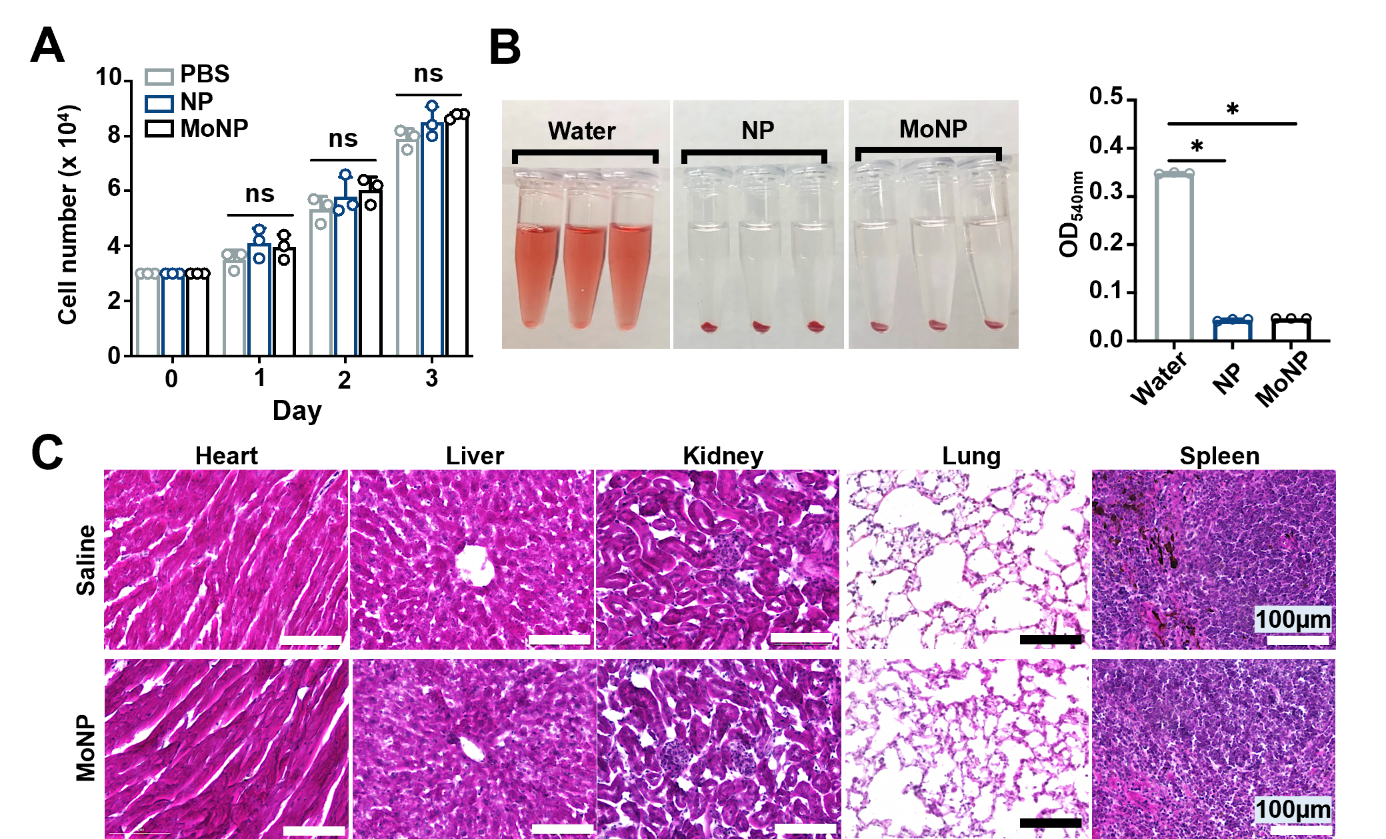

**Figure S5. *In vitro* and *in vivo* biocompatibility assessment of MoNP.** (A) Numbers of live endothelial cells (EC) after incubating with MoNP or NP for 1, 2, and 3 days. n = 3. ns indicates non-significance. (B) Hemocompatibility assay: MoNP or NP resuspended in saline were added to aliquots of mouse whole blood and incubated for 1 hour at 37°C. After low-speed centrifugation, the tubes were imaged, and the absorbance at 540 nm was measured using a plate reader. Deionized water-added blood was used as a control group. *p < 0.05 vs. water. (C) Histological analysis of major organs isolated from the ApoE^-/-^ mice receiving 3 intravenous injections of MoNP or saline.

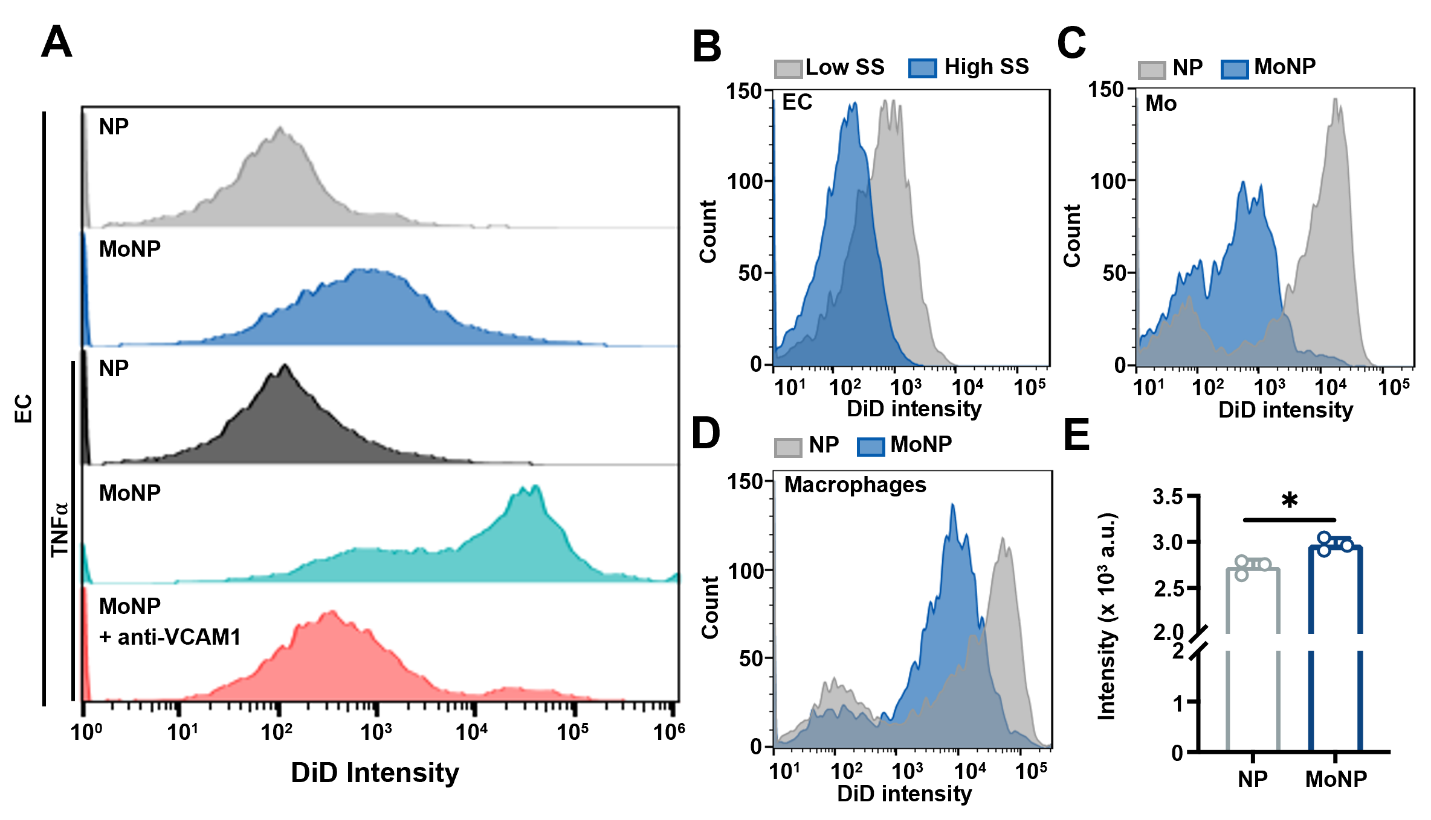

**Figure S6. *In vitro* cellular uptake of MoNP by EC and phagocytes.** (A-D) Flow cytometry results showing the intracellular DiD signal in (A) untreated EC, TNFα-pretreated EC, or TNFα-/anti-VCAM1-pretreated EC, (B) EC under high or low shear stress (SS), (C) Mo, and (D) macrophages. (E) Mouse whole blood was incubated with MoNP-DiD or NP-DiD at 37°C with constant shaking at 120 rpm for 4 hours. After incubation, the samples were centrifuged to collect the plasma, and the fluorescent intensity was measured using a plate reader. n = 3. *p < 0.05 vs. NP-DiD.

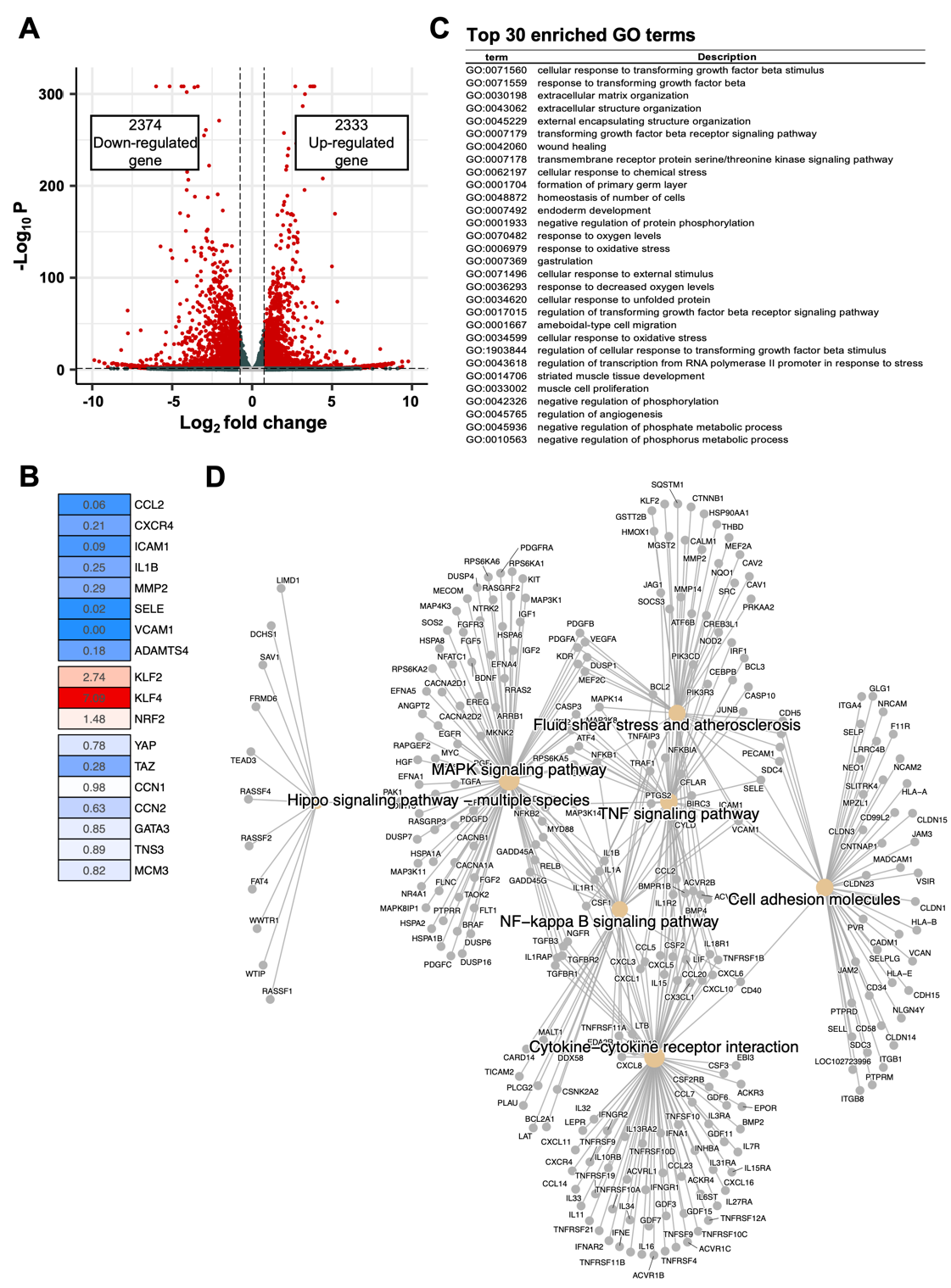

**Figure S7. RNA-seq analysis of inflamed EC under MoNP-VP and MoNP treatments.** (A) The volcano map showed 4,707 differentially expressed genes (DEGs) in TNFα/MoNP-verteporfin (VP)-treated EC vs. TNFα/MoNP-treated EC, including 2,333 upregulated genes and 2,374 downregulated genes. Magenta dots represent genes with |log_2_FoldChange| > 0.75 and p < 0.05. The red nodes represent upregulated and downregulated DEGs. (B) The heatmap of fold change for atheroprone, atheroprotective, and YAP/TAZ-associated genes in TNFα/MoNP-VP-treated EC vs. TNFα/MoNP-treated EC. (C) The top 30 enriched Gene Ontology (GO) terms of DEGs. (D) The Kyoto Encyclopedia of Genes and Genomes (KEGG) pathway enrichment cnetplot showed the enriched pathways of DEGs in response to MoNP-VP vs. MoNP in TNFα-treated EC.

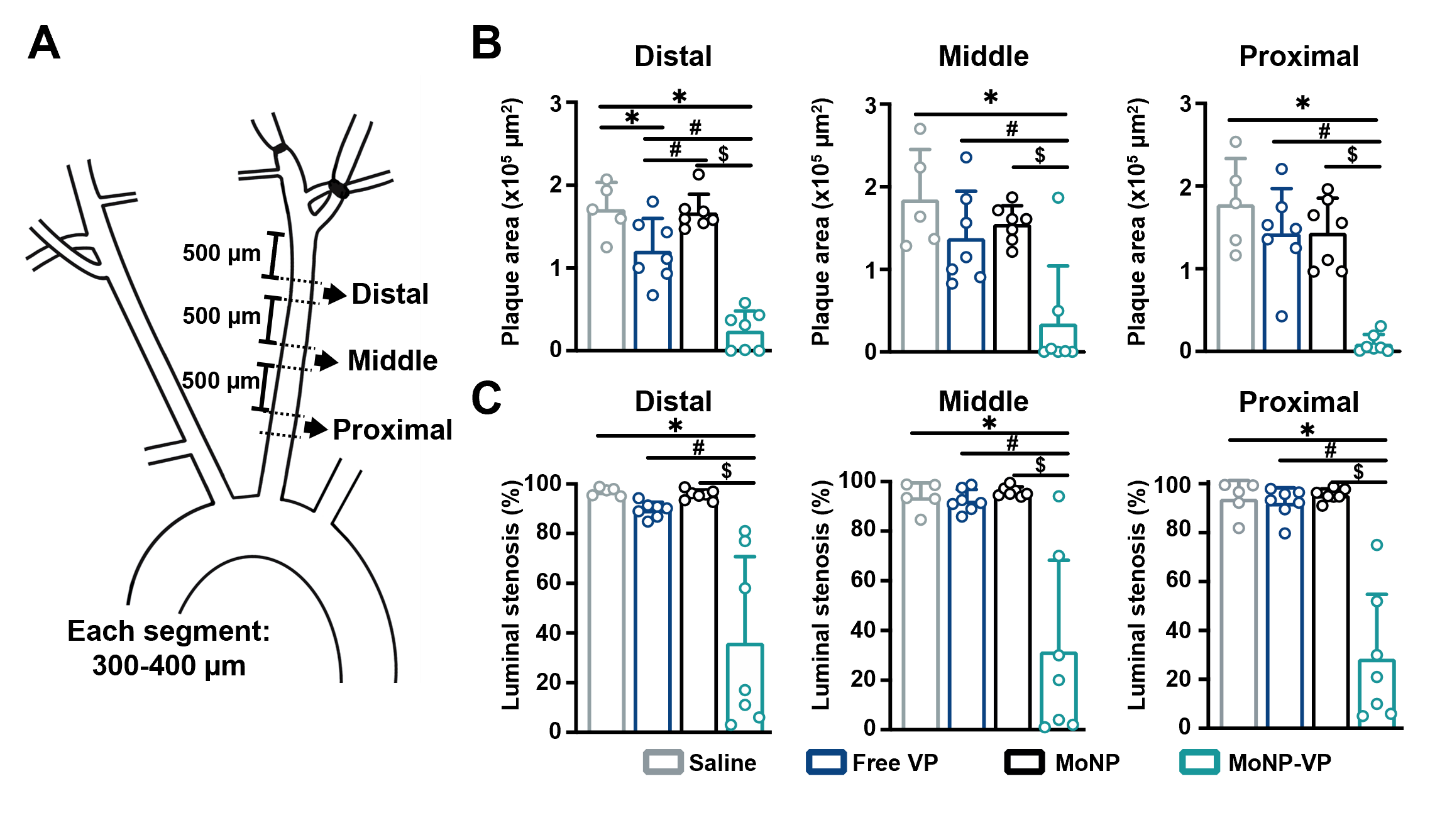

**Figure S8. Quantification of carotid atherosclerosis in ApoE^−/−^ mice under MoNP-VP treatment.** (A) The partially ligated left carotid artery (LCA) was divided into three segments, the distal, middle, and proximal regions, starting from the carotid bifurcation. (B) Quantitative analysis of Oil Red O-positive staining in the distal, middle, and proximal segments of LCA in ApoE^−/−^ mice injected with MoNP-VP, free VP, MoNP, or saline. (B) The Oil Red O-positive area and (C) the degree of luminal stenosis of the LCA were quantified, respectively. n = 7 each for MoNP-VP, free VP, and MoNP, and n = 5. *p < 0.05 vs. saline, ^#^p < 0.05 vs. free VP, and ^$^p < 0.05 vs. MoNP.
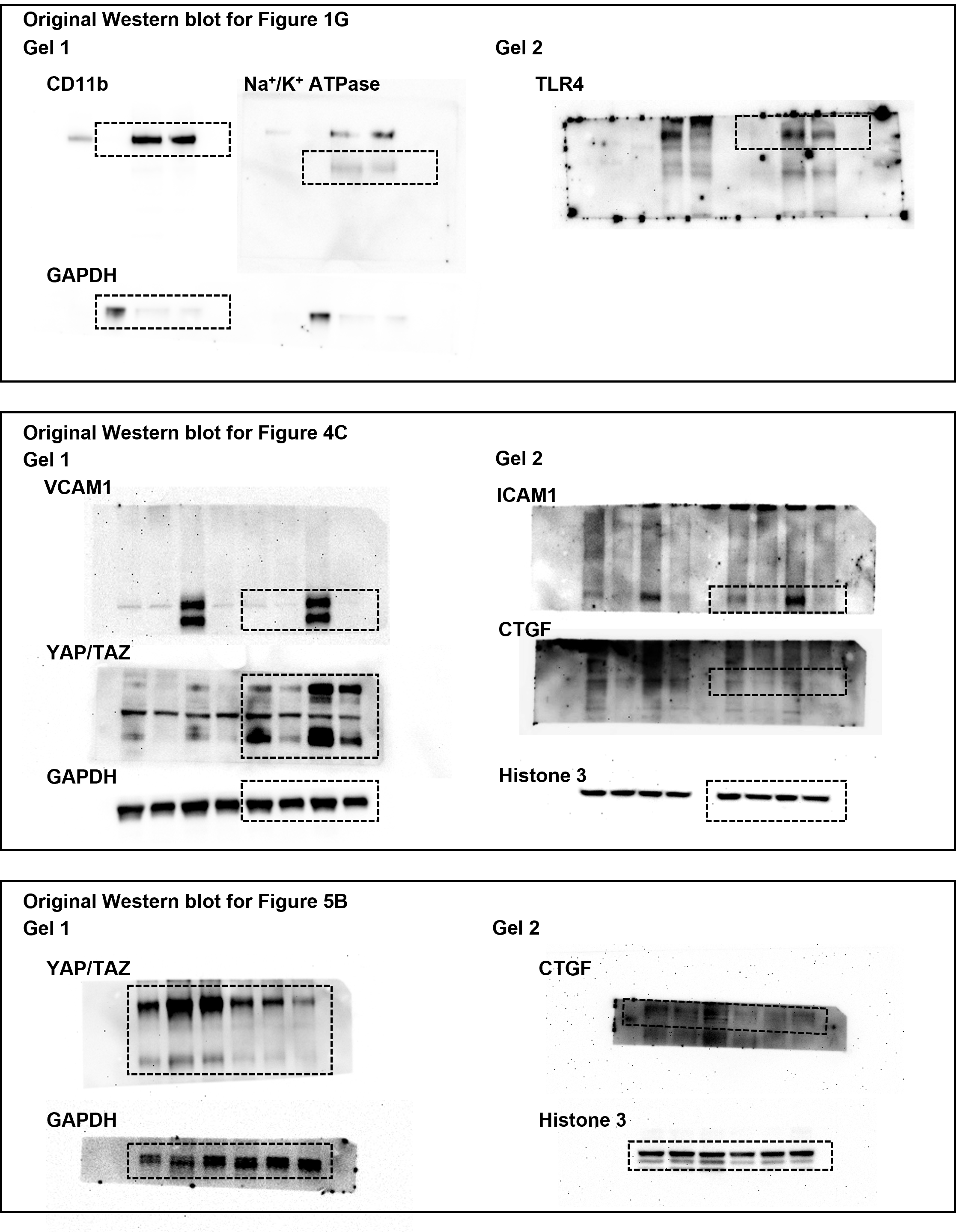

**Figure S9. Uncropped Western blot images**

**Table S1. The metabolic panel of mice receiving MoNP-VP**

| **Tests** | **MoNP-VP (n=6)** | **Reference values [1]** |
| --- | --- | --- |
| Albumin (g/dL) | 3.48 ± 0.13 | 2.6-5.4 |
| Alkaline phosphatase (U/L) | 84 ± 8 | 16-200 |
| Alanine aminotransferase (U/L) | 59.83 ± 30.09 | 22-133 |
| Total bilirubin (mg/dL) | 0.56 ± 0.15 | 0.1-0.9* |
| Blood urea nitrogen (mg/dL) | 21.33 ± 2.66 | 2-71 |
| Calcium (mg/dL) | 10.20 ± 0.67 | 6.8-11.9* |
| Phosphorus (mg/dL) | 6.2 ±1.81 | 6.0-11.3* |
| Creatinine (mg/dL) | 0.2 ± 0.0 | 0.1-1.8 |
| Glucose (mg/dL) | 192.33 ±70.54 | 60-133 |
| Sodium (nmol/L) | 141.33 ± 2.34 | 153-175 |
| Potassium (nmol/L) | 6.38 ± 1.62 | 6.5-9.7* |
| Total Protein (g/dL) | 4.67 ± 0.19 | 4.6-7.3 |
| *The reference values for CD1 were utilized in cases where the corresponding values for C57BL6 are unavailable. | | |

**Table S2. List of primers for qRT-PCR (F: forward; R: reverse)**

| **Gene** | **Species** | **Sequence (5’ 🡪 3’)** |
| --- | --- | --- |
| VCAM1 | human | F: ggcagagtacgcaaacactt R: acaggattttcggagcagga |
| ICAM1 | human | F: cttgagggcacctacctctg R: cattatgactgcggctgcta |
| GAPDH | human | F: gcaccaccaactgcttagc R: atgatgttctggagagcccc |
| CTGF | human | F: ggcccagacccaactatgat  R: aggaggcgttgtcattggta |

**Reference**

[1]	F.W. Quimby, R.H. Luong, Clinical Chemistry of the Laboratory Mouse, The Mouse in Biomedical Research: History, Wild Mice, and Genetics: Volume 1-4, Second Edition. 1–4 (2007) 171–216. https://doi.org/10.1016/B978-012369454-6/50060-1.
